# Multi-state structure determination and dynamics analysis reveals a new ubiquitin-recognition mechanism in yeast ubiquitin C-terminal hydrolase

**DOI:** 10.1101/2021.04.22.440356

**Authors:** Mayu Okada, Yutaka Tateishi, Eri Nojiri, Tsutomu Mikawa, Sundaresan Rajesh, Hiroki Ogasa, Takumi Ueda, Hiromasa Yagi, Toshiyuki Kohno, Takanori Kigawa, Ichio Shimada, Peter Güntert, Yutaka Ito, Teppei Ikeya

## Abstract

Despite accumulating evidence that protein dynamics is indispensable for understanding the structural basis of biological activities, it remains challenging to visualize the spatial description of the dynamics and to associate transient conformations with their molecular functions. We have developed a new NMR protein structure determination method for the inference of multi-state conformations using multiple types of NMR data, including paramagnetic NMR and residual dipolar couplings, as well as conventional NOEs. Integration of these data in the structure calculation permits delineating accurate ensemble structures of biomacromolecules. Applying the method to the protein yeast ubiquitin hydrolase 1 (YUH1), we find large dynamics of its N-terminus and crossover loop surrounding the active site for ubiquitin-recognition and proteolysis. The N-terminus gets into and out of the crossover loop, suggesting their underlying functional significance. Our results, including those from biochemical analysis, show that large motion surrounding the active site contributes strongly to the efficiency of the enzymatic activity.

## Introduction

It is well known that the dynamics of three-dimensional (3D) protein structures enables multiplicity and flexibility of molecular functions. Numerous experimental reports have demonstrated that the functional or binding sites of proteins are often not rigid but rather flexible, allowing multiple roles or adaptive interactions with several other molecules^1^. Intrinsically disordered proteins (IDPs), for example, possess extraordinarily large structural flexibility that confers them unique properties to interact with several other counterparts^2^. Considering that flexible protein structures are susceptible to their surrounding environment, it is crucial to study the molecular motions and 3D structures of proteins at atomic resolution under near-physiological conditions. Nuclear Magnetic Resonance (NMR) spectroscopy is currently the only technique to investigate the dynamics and 3D conformations of proteins in solution or even in living systems^3,4^. Data from solution NMR includes information on the protein behavior and dynamics as a structural ensemble, and hence various NMR methodologies have been proposed for the extraction of their physical properties from the spectra. For instance, NMR experiments for the quantification of protein dynamics, such as CPMG relaxation dispersion^5^, DEST^6^, and CEST^7^, provide particularly useful information on molecular motion at multiple timescales and lowly populated “excited-state” conformations. However, it remains a challenge to visualize dynamic protein structural ensembles and to elucidate their correlations with molecular functions. To date, several studies have addressed the reconstruction of structural ensembles by extracting dynamical 3D structural information from residual dipolar couplings (RDCs) ^8^, paramagnetic relaxation enhancements (PREs)^9,10^, solvent PREs^11^, and exact nuclear Overhauser effects (eNOEs) ^12^. However, many of these studies investigated disordered proteins, did not perform *de novo* protein structure determination exclusively from experimental data, or were applied to small proteins using only one or two types of NMR data. The NOE-type experiments provide distance information from a large number of hydrogen pairs in a molecule, which is necessary to obtain the complete structure of a protein containing detailed side-chain coordinates, but cover only short distances up to 5–6 Å. On the other hand, monitoring large dynamic conformations with high reliability also requires structural information for longer distances. Paramagnetic effects, PREs and pseudcontact shifts (PCSs), provide long-range distance and angle information for atoms located up to 40 Å from the paramagnetic center^13^. The determination of the complete structure with high precision exclusively from paramagnetic experiments, however, would require an excessive number of experiments to conjugate paramagnetic tags at different positions on the molecular surface to collect structural information covering the whole molecule. In practice, it is thus necessary for *de novo* structure determination to integrate multiple NMR data including NOEs, chemical shifts, PREs, PCSs, and RDCs.

Meanwhile, the visualization of the conformational distribution in the presence of large motions requires an ensemble structure calculation considering multiple states rather than a single conformation. *De novo* multi-state structure determination does not need prior knowledge of 3D information, and permits finding underlying protein dynamics and functions. Hence, we employed the multi-state structure calculation method implemented into the program CYANA, which was originally applied to eNOE data of the protein GB1^12^. Based on the original algorithm, we here expand this method to other long-range experimental data, in particular for handling PCS data. The PCS *Δ*χ tensors are determined iteratively based on intermediate structure ensembles during the calculation. To salvage NOEs that were not found by the conventional method due to large dynamics, the automated NOE assignment^14^ was also performed repeatedly each time when the ensemble structures were updated. For systems with a large conformational distribution, these improvements for the multi-state calculation are expected to be particularly effective.

As a model system for the multi-state structure determination with multiple NMR data sets, we selected yeast ubiquitin hydrolase 1 (YUH1), which is composed of 236 residues with a molecular mass of 26.4 kDa. It belongs to the ubiquitin C-terminal hydrolases (UCHs) family with a function to detach small peptides or chemical adducts from the C-terminus of ubiquitin^15,16^. In many enzymes associating with ubiquitination and deubiquitination, UCHs mainly contribute to the maintenance of the homeostatic equilibrium of ubiquitin level in eukaryotic cells by recycling ubiquitin after the proteasomal degradation in cell systems^17^. Since the primary sequence of ubiquitin is highly conserved in eukaryotes from yeast to human, the enzymatic processes of ubiquitination and deubiquitination must be similar. Despite numerous studies for UCHs, however, the molecular mechanism of ubiquitin recognition by UCH remains a fundamental question in molecular biology. The crystal structure of the YUH1 complex with ubiquitin aldehyde (Ubal)^18^ demonstrates that the binding surface is divided primarily into two regions, one surrounding the enzymatic active site and the other interacting with the globular domain of ubiquitin (Supplementary Fig. 1a). Ubiquitin comprises a globular domain and a flexible C-terminus that anchors to other proteins during ubiquitination. The interaction surface of YUH1 with the globular domain of ubiquitin is presumably needed for the enzymatic reaction by tightly holding the ubiquitin body in place. It is known that the active site of YUH1 consists of the cysteine nucleophile (Cys90), an adjacent histidine (His166), and an aspartate (Asp181) ^18^. The surrounding area of the active site contains a loop which is referred to as the crossover loop (147–163) and the N-terminus (1–14) of YUH1, forming a hole where the C-terminus of Ubal is entirely embedded, probably for the suppression of its flexibility (Supplementary Fig. 1a). In there, the ubiquitin C-terminus and the YUH1 N-terminus pass in parallel through the crossover loop, resulting in a tight packing of the hole. This observation raises the additional question as to how YUH1 efficiently and quickly incorporates the C-terminus of ubiquitin into the deep hole, catalyzes the hydrolysis at the optimal position, and releases it for the next cycle of the reaction. The configuration of the active site is well conserved among the UCH family^19-22^, indicating that this unique structural feature is indispensable for the enzymatic activity. Navarro *et al*. propose that the crossover loop restricts or filters the size of adducts of ubiquitin^23^. In the structural studies of free and complex ubiquitin carboxyl-terminal hydrolase isozyme L3 (UCH-L3) by X-ray crystallography, the atomic coordinates in the crossover loop were clearly determined in the complex but not in the monomer^24^, suggesting that the crossover loop is fairly flexible in the free state. This flexibility probably enables the efficient insertion of the ubiquitin C-terminus into the deep hole at the active site. Indeed, in our previous solution NMR study of free YUH1, the ^1^H-^15^N
 correlation cross-peaks in the 2D HSQC spectrum were distinctly strong for this region, and very few medium/long-range NOEs were detected in the ^15^N-separated NOESY spectrum^25^. On the other hand, temperature factors are relatively low for this region in the crystal structure of the complex YUH1 with Ubal, implying that the crossover loop gets rigid by the interaction with ubiquitin (Supplementary Fig. 1b). Considering that the flexible property of YUH1 is presumably susceptible to the surrounding environment as described above, it is essential to elucidate its dynamics and structural distribution in the solution state.

In this article, we investigate the 3D structure of free YUH1 and its dynamics in solution by multi-state structure determination using multiple NMR data sets. Visualizing the ensemble structures of YUH1 by this method demonstrates extraordinarily large dynamics not only in the crossover loop but also in the N-terminus of YUH1. The latter raises a further question as to whether these dynamics correlate with a molecular function. Hence, based on the conformational distribution and functional evidence, we propose an additional role of the N-terminus (henceforth referred to as ‘the gating loop’ due to its location and the addition role elucidated in this study) and discuss the molecular mechanism of ubiquitin hydrolase by YUH1.

## Results

### Protein structure and dynamics analysis by conventional approaches

The backbone ^1^H^N, 15^N, ^13^C^α, 13^C′ and sidechain ^13^C^β^ resonances of free YUH1 have already been assigned in our previous study^25^. By analyzing 3D triple-resonance NMR spectra as well as 3D ^15^N- and ^13^C-separated NOESY spectra, we now assigned approximately 95% of H^α^, and 98% of H^β^ resonances of YUH1 (Fig. 1a, and Supplementary Fig. 2). Methyl resonances were first classified into Ala, Ile, Leu, and Val by amino acid-selective labeling, and then 99% of the methyl ^1^H-^13^C resonances were assigned by analyzing 3D HC(C)H-TOCSY, ^15^N- and ^13^C-separated NOESY spectra (Supplementary Fig. 3c). Similarly, amino-acid type classification for aromatic residues was performed by amino acid-selective labelling, and 89 out of 104 (86%) of the aromatic ^1^H-^13^C resonances were assigned by analyzing 3D ^15^N- and ^13^C-separated NOESY spectra (Supplementary Fig. 3d). Using manual and automated NOE assignment implemented into the program CYANA^14^, overall 4798 distance restraints (including 1917 long-range) involving methyl and aromatic groups were assigned and used in the structure calculation. In addition, 319 dihedral angle restraints for ϕ and ψ angles were derived from backbone chemical shifts and restraints for 29 hydrogen bonds were collected by scalar-couplings (^3h^*J*_N-C’_) from a long-range HNCO experiment^26^.

**Fig. 1.**
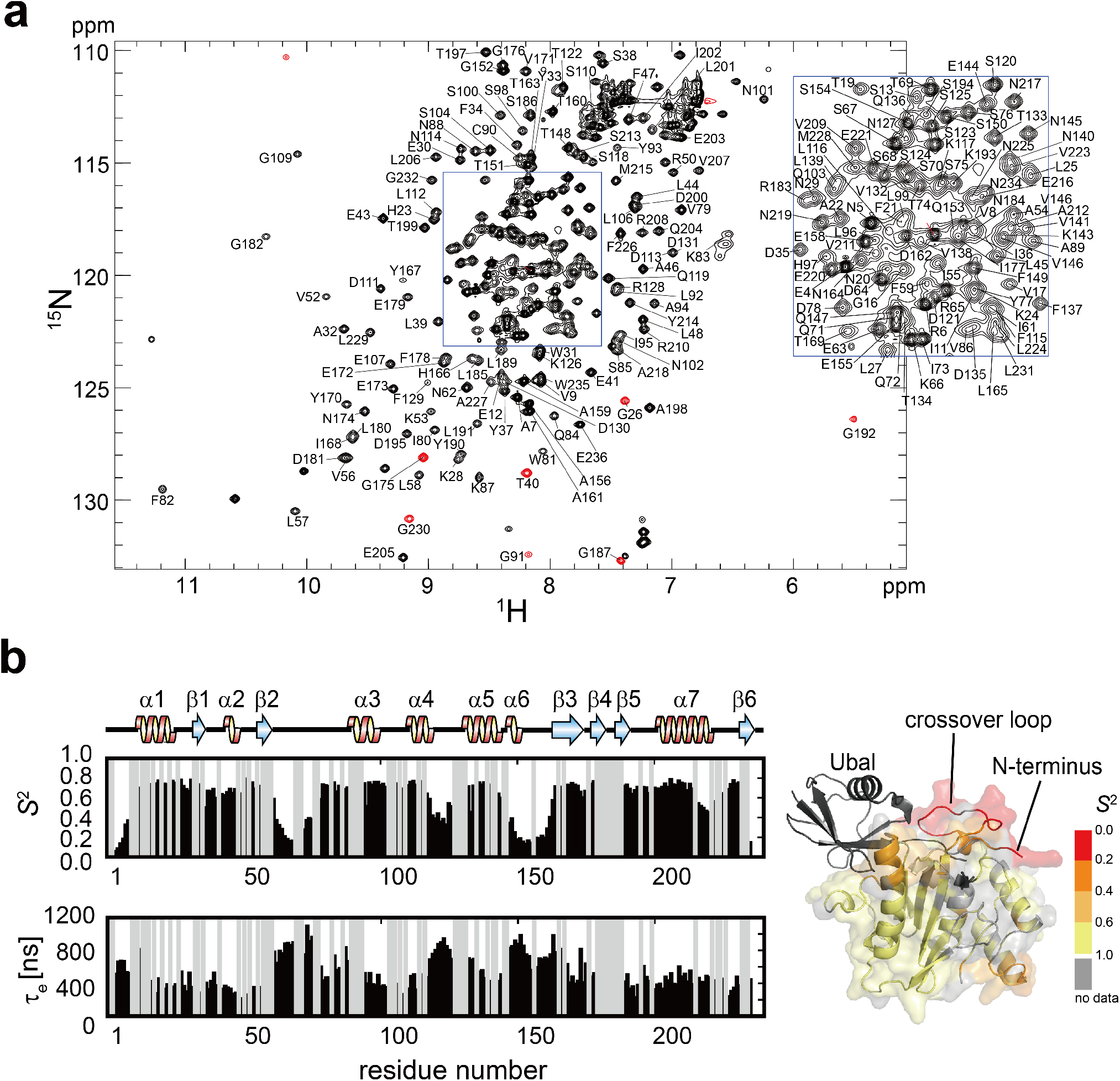
Backbone resonance assignments and dynamics of YUH1. **a** 2D ^1^H-^15^N HSQC spectrum and backbone resonance assignments of YUH1. The center area surrounded by the blue frame is separately shown on the right top. Positive and negative (aliased on the ^15^N axis) signals are color-coded in black and red, respectively. **b** Generalized order parameters (*S*^2^) and effective rotational correlation times (*τ*_e_) for the ^15^N-^1^H vector of each residue calculated from *T*_1_/*T*_2_ relaxation and heteronuclear {^1^H}-^15^N NOE measurements. The *S*^2^ values are mapped on the crystal structure of the YUH1-Ubal complex.

We first performed a conventional structure calculation exclusively using distance and dihedral angle restraints from NOEs, hydrogen bonds, and chemical shifts. The resulting ensemble of 20 structures was well converged with an average backbone RMSD of 0.56 Å to the mean coordinates except for the N-terminus (gating loop; 1–14), the crossover loop (L9: 149–164), and the “invisible” region in the YUH1-Ubal crystal structure (L5: 63–77) (Supplementary Fig. 4a and Supplementary Table. 1). Except for these three regions, the global fold of free YUH1 was similar to that of YUH1 in the YUH1-Ubal crystal structure with a backbone RMSD of 2.73 Å (Supplementary Fig. 4c). In contrast, the location of the gating loop was significantly different. In the YUH1-Ubal co-crystal, the gating loop passes through the crossover loop and lies besides the active center in the complex structure (Supplementary Fig. 4b), whereas it is oriented in the opposite direction pointing away from the crossover loop in the solution structure (Supplementary Fig. 4a, henceforth this conformation is referred to as the ‘open conformation’).

The number of long-range NOEs in the abovementioned three regions (gating loop, L5, and L9) is lower than elsewhere (Supplementary Fig. 5), suggesting large dynamics of these regions. Next, to quantify the dynamics, we measured the ^15^N longitudinal (*T*_1_) and transverse (*T*_2_) relaxation times, and the steady-state heteronuclear {^1^H}-^15^N NOE (Supplementary Fig. 6). From these, the overall rotational correlation time *τ*_c_, and effective correlation times *τ*_e_ and generalized order parameters *S*^2^ for each ^15^N-^1^H vector were estimated by a Lipari-Szabo model-free analysis from the relaxation experiments (Fig. 1b). The *τ*_c_ from the model-free analysis was 18.5 ns, which is in good agreement with a rotational correlation time of 18.0 ns calculated by the Stokes-Einstein-Debye equation, validating this model-free analysis of the NMR relaxation data. The *S*^2^ order parameters for the gating loop, L5, and L9 regions were below 0.6, demonstrating that they indeed possess large dynamics, while their temperature factors in the crystal structure of the complex in were relatively low (Supplementary Fig. 1b). Interestingly, reports on free UCH-L3, which have approximately 60% sequence homology with YUH1 (Supplementary Fig. 7), have discussed that the crossover loop (L5) is quite flexible in the free state for efficient incorporation of the ubiquitin C-terminus into the active site^20,27^. However, these discussions were simply based on the fact that the crossover loop is invisible in the electron density of free UCH-L3. Consequently, our finding on YUH1 in solution is the first direct evidence for the large dynamics of the crossover loop.

Although ^15^N-relaxation analysis reveals the dynamics of the gating loop, L5, and L9 regions of YUH1, the conformations of these regions found in the current single-state solution structure calculations were somewhat uncertain. The conformation of the gating loop was extraordinarily different from that of the complex in the crystal (Supplementary Fig. 4). As mentioned above, in the YUH1-Ubal crystal structure the N-terminal chain passes through the crossover loop and is located close to the active site Cys90. Comparing this with the gating loop location in the crystal structure of free UCH-L3, Johnston *et al*. have proposed that the specificity of UCH to ubiquitin is enhanced by the steric obstruction of its N-terminus (gating loop) on the enzymatic active site^18^. Indeed, a sidechain of the UCH-L3 N-terminus (gating loop) completely buries the catalytic cysteine nucleophile in the free structure. Since the current solution structure did not support this model, we initiated PRE, PCS, and RDC experiments to obtain long-range structural information in order to obtain detailed information on the relative orientation of the crossover loop and gating loop against the globular domain of YUH1 in solution.

### PCS, PRE, RDC experiments for long-range structural information

To conjugate paramagnetic tags to different sites on the protein surface, we produced four separate cysteine mutants. In all these mutants, the wild-type cysteine 90 was replaced with serine (C90S), and then one residue was replaced by a cysteine, N5C, S104C, N140C, and N225C (Supplementary Fig. 8). We observed no large perturbations of the ^1^H-^15^N-HSQC spectra between the wild type YUH1 and the four mutants except for the peaks of the replaced residues (Supplementary Fig. 9), indicating that the mutations did not alter the original fold of YUH1.

For PCS measurements, DOTA-M8-SPy^28^ and DO3MA-3BrPy^29^ were used depending on the reaction efficiencies at the cysteine mutation sites. Strong PCSs were observed in ^1^H-^15^N-HSQC spectra of the N140C mutant combined with DOTA-M8-SPy and DO3MA-3BrPy (Fig. 2a and b), indicating sufficiently suppressed mobility of the tags and thereby accurate structural information. The paramagnetic Dy^3+^ ion yielded shifted single ^1^H-^15^N correlation cross peaks for most of the affected residues, demonstrating that multiple orientations of the paramagnetic tag itself did not occur in this experiment. In contrast, surprisingly, cross peaks mostly corresponding to the gating loop and the crossover loop were split into two or three. For instance, in the ^1^H-^15^N HSQC spectrum of the N140C mutant combined with [Dy^3+^ (DOTA-M8-SPy)], Gly3 and Val9 showed cross peaks split into three, and Ala7, Ala156, Ala159, and Ala161 yielded cross peaks split into two. Figure 2b compares observed PCS values with those predicted from the YUH1 coordinates in the YUH1-Ubal co-crystal. The PCSs for the residues in the gating loop and the crossover loop are significantly different from the predicted values, whereas those for the rest of the residues show a good match. These results indicate that the gating loop and the crossover loop possess multiple conformations, each of which is different from that in the YUH1-Ubal crystal. Exchange between these conformations is slow on the NMR time-scale, suggesting the presence of ∼msec dynamics.

**Fig. 2.**
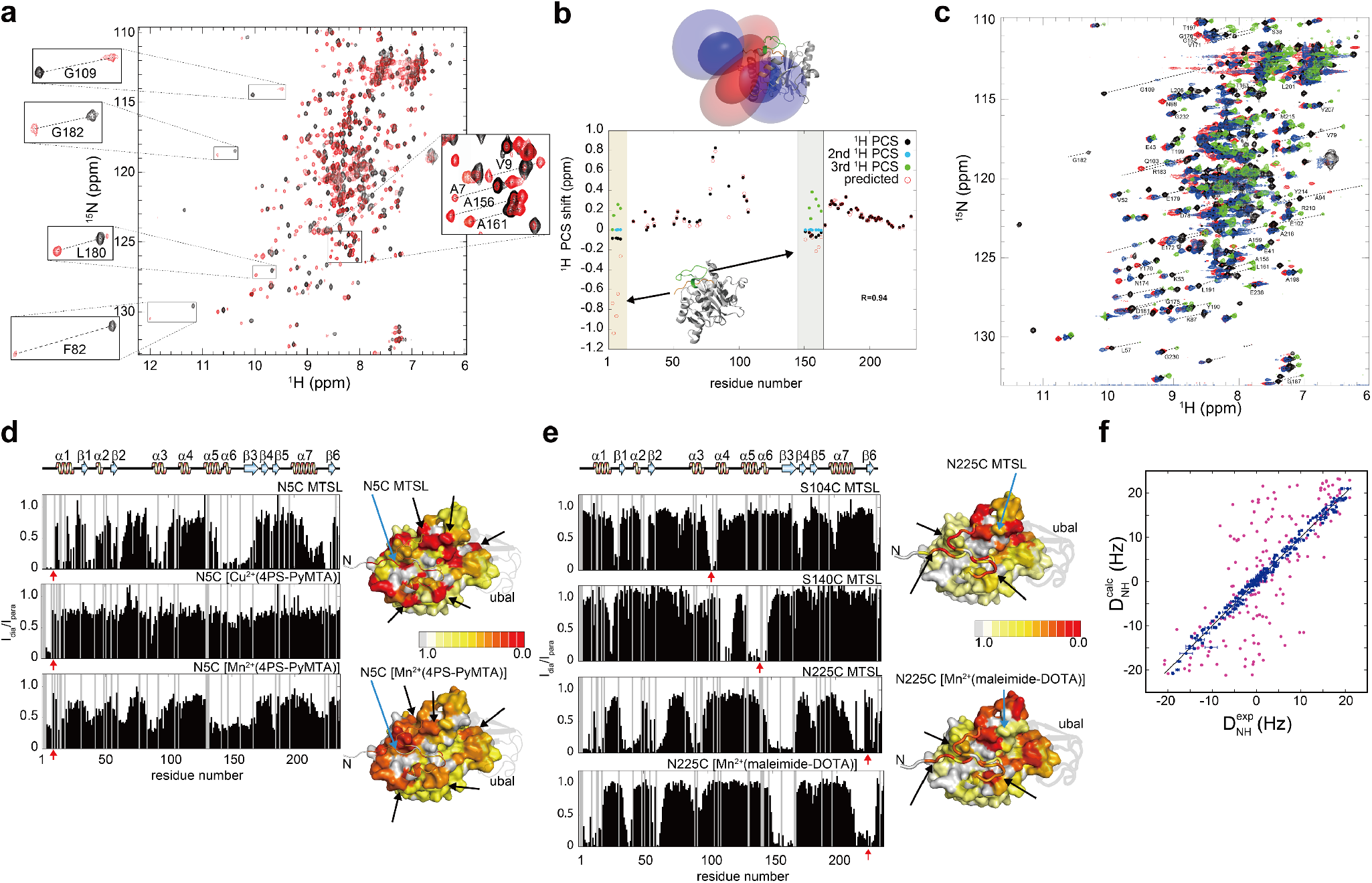
PRE, PCS and RDC measurements on YUH1. **a** ^1^H-^15^N HSQC spectra of YUH1(N140C) tagged with diamagnetic [Lu^3+^(DOTA-M8-SPy)] (black) and paramagnetic [Dy^3+^(DOTA-M8-SPy)] (red). Several representative cross peaks are shown in boxes. **b** Comparison of experimental PCSs from DOTA-M8-SPy for the free YUH1(N140C) with those predicted from the YUH1 coordinates in the YUH1-Ubal crystal structure (PDB 1CMX). The regions corresponding the gating loop (1–14) and crossover loop (L5) are shown in orange and light green strips, respectively. The *Δ*χ tensors are represented by PCS isosurfaces, with positive and negative shifts shown in blue and red, respectively. **c** ^1^H-^15^N HSQC spectra of YUH1(N140C) tagged with DO3MA-3BrPy in the presence of diamagnetic Lu^3+^(black) and paramagnetic Dy^3+^(red), Tb^3+^ (blue), and Tm^3+^ (green). The crowding area and low intensity peaks are separately shown in Supplemental Figure 10. **d** Ratios of peak heights between paramagnetic and diamagnetic samples (*I*_para_/*I*_dia_) of the free YUH1(N5C) tagged with MTSL, [Cu^2+^(4PS-PyMTA)], and [Mn^2+^(4PS-PyMTA)]. Red arrows indicate the spin-labelled positions. Residues are color-coded by *I*_para_/*I*_dia_ ratios, as indicated in the color bar, on ribbon and surface models of the YUH1-Ubal crystal structure. The conformation for residues 1– that lack coordinates in the crystal structure was modeled by the CYANA regularize module^50^. Residues without data and Ubal are color-coded in gray and black, respectively. The black arrows indicate the regions that are far from the paramagnetic center but show strong PREs. **e** Ratios of peak heights between paramagnetic and diamagnetic samples (*I*_para_/*I*_dia_) of YUH1 S104C tagged with MTSL, S140C with MTSL, N225C with MTSL, and N225C with [Mn^2+^(maleimide-DOTA)], color-coded as in panel d. **f** Correlation between observed RDCs and RDCs calculated on the basis of YUH1 structures computed by the conventional single-state method (red) or multi-state structure calculation (blue). All spectra were recorded at 303 K and a magnetic field strength of 14.1 T.

To validate the split PCSs, we repeated the experiments by employing DO3MA-3BrPy with several lanthanoid ions, Dy^3+^, Tb^3+^, and Tm^3+^, which are expected to provide various PCS effects due to their different paramagnetic properties. As expected, the resulting spectra exhibited similar doubled or tripled peaks but with different PCS values, again mostly for the residues in the gating loop and the crossover loop (Fig. 2c and Supplementary Fig. 10).

For PRE measurements, 4PS-PyMTA^30^, maleimide-DOTA^31^, and MTSL^32^ were used as the paramagnetic probes by conjugating them to substituted cysteine residues through thioether or disulfide bonds. Strong PREs were observed for the N5C, S104C, N140C, and N225C mutants conjugated with MTSL, N225C with [Mn^2+^(maleimide-DOTA)], and N5C with 4PS-PyMTA. We expected that the PRE-experiments on the N5C mutant, in which the paramagnetic tag is conjugated directly to the gating loop would validate or refine the gating loop conformations in solution. However, surprisingly, strong PREs were observed not only in the region surrounding the gating loop in the YUH1-Ubal crystal structure but also far away from it (Fig. 2d). For instance, strong PREs by the MTSL and [Cu^2+^(4PS-PyMTA)] tags at N5C were observed in the region 13–18, 35–42, 60–64, 97–99, 152–155, 164–169, and 224–227 which are more than 20 Å distant from Cys9 in the crystal structure. This result indicates that the gating loop does not always stay at one particular site seen in the crystal structure but instead moves dynamically to different positions. Note that the spatial motion of the gating loop must be large-scale, since the gating loop first needs to exit from the tunnel formed by the crossover loop in order to approach the regions of residues 13–18, 35–42, 60–64, and 224–227. Similar behavior was also found in the PRE-experiment with MTSL-attached N225C (Fig. 2e). In the complex structure, Arg6 in the gating loop is located over 25 Å apart from Asn225, and Glu4 (missing its coordinates in the complex) would be even further away than Arg6. Nonetheless, strong PREs were observed for residue 4–8. These results also suggest that the gating loop does not stay at the position in the YUH1-Ubal crystal structure but approaches the site of Asn225 for a certain period.

N-H^N^ RDCs were obtained by aligning with phage pf1 and measuring 2D ^15^N-^1^H IPAP HSQC^33^ for the isotropic and aligned samples. We first calculated the alignment tensor and predicted RDCs based on the YUH1 coordinates in the YUH1-Ubal co-crystal. Just as for the results of the PCSs and PREs, the experimental data was not entirely consistent with those from the complex (Fig. 2f), which further supports the hypothesis that the solution structure in the free state is quite different from the one in the YUH1-Ubal crystal.

### Multi-state structure calculation

The data from PCSs, PREs, and RDCs for free YUH1 in solution was not compatible with a single-conformation model, particularly for the gating loop and the crossover loop due to their large dynamic motions. Thus, we adopted multi-state structure calculation using the program CYANA to analyze simultaneously several NMR data sets, including NOEs, chemical shifts, PREs, PCSs, and RDCs. Based on the original algorithm^12^, we expanded this method for handling other long-range experimental data, in particular PCS data. The PCS *Δ*χ tensors were determined iteratively based on an intermediate structure ensemble during the calculation. Automated NOE assignment using network-anchoring and constraint combination^14^ was also performed repeatedly referring to the intermediate multi-state ensemble conformations, so that additional NOEs were collected mostly for the regions with high mobility such as the gating loop and the crossover loop (Supplementary Fig. 11). A cross-validation test determined the final number of the conformational states. The test shows that increasing the number of conformations decreases the CYANA target function until a plateau is reached with a five-state model, indicating that the five-state model fulfills best the experimental data (Fig. 3a). Compared with the single-state structures from the conventional method, the five-state ensemble conformations are in good agreement with the RDC data. The correlation between the back-calculated and experimental RDCs improved dramatically over the single-state structure (Fig 2f).

**Fig. 3.**
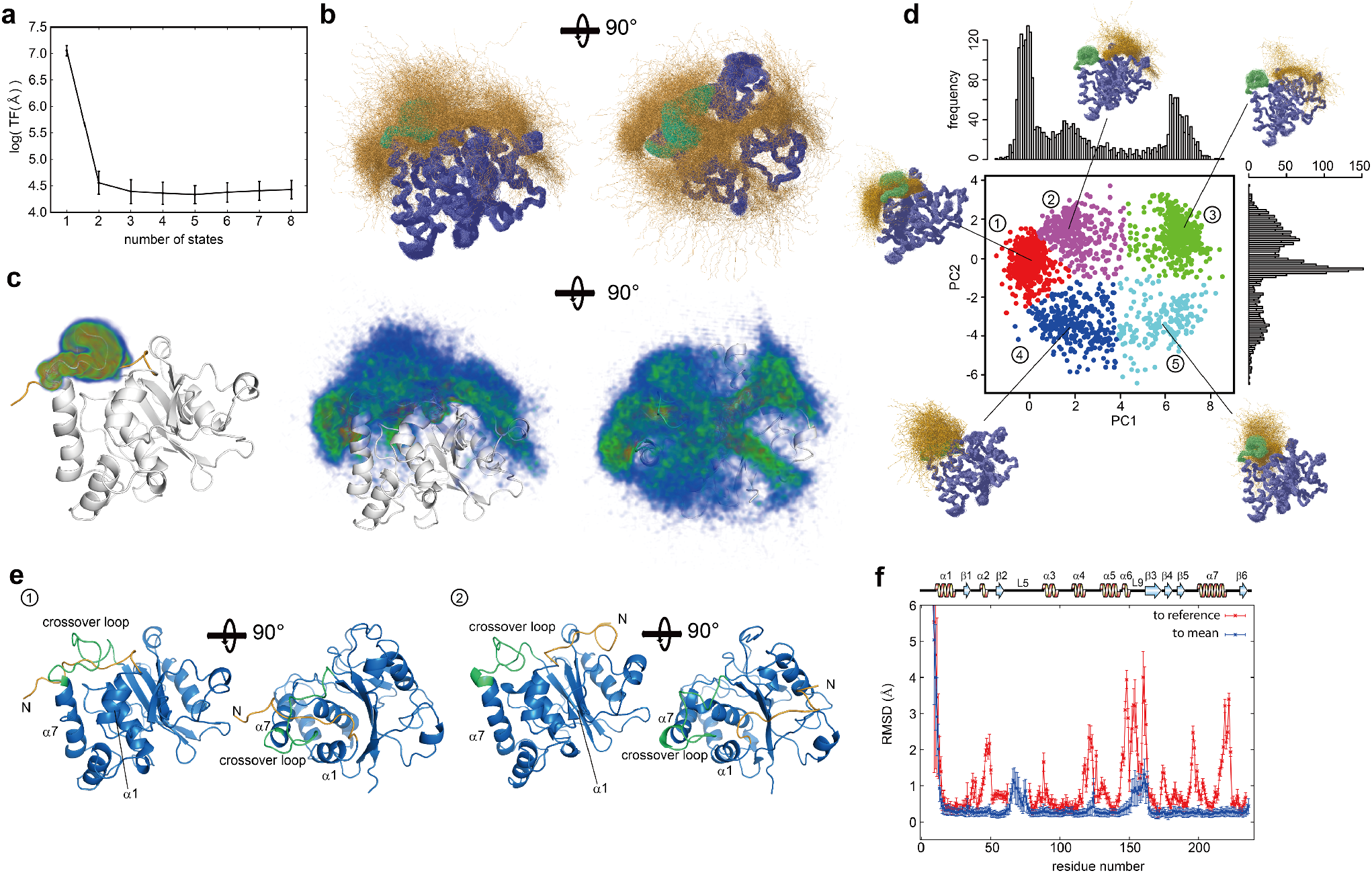
Ensemble structures of free YUH1. **a** Cross-validation tests highlighted by the CYANA target function which is the (weighted) sum of the squared violations of all conformational restraints. **b** 2500 conformers obtained by multi-state structure calculation with a five-state model, showing the backbone atoms N, C^α^, C’. The gating loop and the crossover loop are colored in orange and green, respectively. **c** Atomic probability density maps showing the conformational space sampled by this method for the crossover loop (left) and gating loop (center and right). **d** Distribution of 2500 conformers analyzed by principal component analysis (PCA). The horizontal and vertical axes show the first and second principal components, PC1 and PC2. The plot is classified by k-means clustering shown in different colors. Clusters are numbered in the order of their size. The histograms along the horizontal and vertical axes indicate the distributions on PC1 and PC2. Ribbon models are representative conformations at the cluster centers. **e** Representative conformations at the centers of the largest and second largest clusters in d, color-coded as in b. **f** RMSD per residue and its standard deviation for the 2500 conformations, computed for C^α^ to the mean structure (blue) and the crystal structure of YUH1-Ubal (red).

Except for the gating loop, the crossover loop (L9), and invisible regions in the YUH1-Ubal crystal structure (L5), the global fold of the five-state ensemble structures was very similar to the complex with a backbone RMSD of 2.02 Å for residues 15–62, 78–148, and 165–236 (Fig. 3b-d and 4a). Meanwhile, the gating loop and crossover loop spread out over an extensive area in conformational space, but their structures are not entirely random. To analyze the structural distribution based on representative variances of 3D coordinates, we performed principal component analysis (PCA) and k-means clustering with the assumption of the existence of five clusters in the plane spanned by the first and second principal components (PC1 and PC2) (Fig. 3d). The largest cluster was the most compact distribution on the PC1 and PC2 plane. 1059 out of all 2500 conformers (∼42%) belong to this cluster and are very similar to the YUH1-Ubal crystal structure with an average backbone RMSD of 2.26 Å for all residues present in the crystal structure, 6–62 and 78–234. The structures of the first cluster are in the closed form in which the gating loop passes through the crossover loop and the loop orients toward the active center’s side (Fig. 3e, left panel). This suggests that the two regions are extraordinarily flexible but still mostly exist as the closed-form even in the free state. The other clusters except the fifth comprised conformers in which the gating loop does not pass through the crossover loop, and is oriented in different directions against the crossover loop. The second-largest cluster (∼23%) mostly comprises conformations in which the gating loop is located on the binding site for the ubiquitin globular domain (Fig. 3e). The binding site constitutes a large groove, into which the gating loop fits well. The third and fourth cluster are roughly in between the first and second, in that the gating loop is rotated approximately 90 degrees away from its positions in the first and second clusters. They might be intermediate conformations in the equilibrium between the first (closed) and second (open) forms. The structures in the fifth cluster contained similar gating loop orientations as in the YUH1-Ubal crystal structure but consisted of two different forms where the gating loop passes over and under the crossover loop, respectively.

**Fig. 4.**
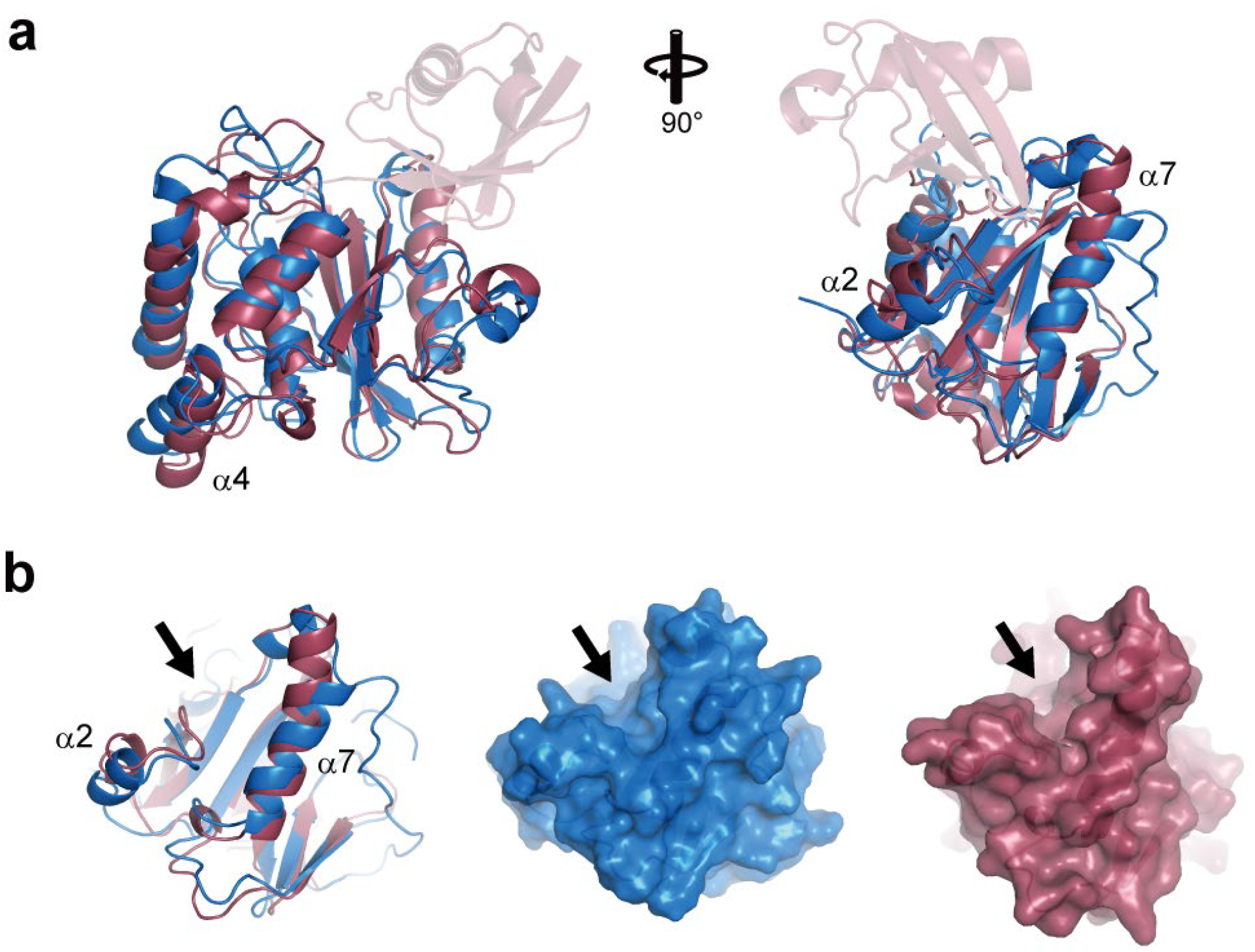
Comparison of the free YUH1 structure with the YUH1-Ubal complex. **a** Superposition of the free YUH1 structure at the center of the largest cluster (light blue), and the complex (dark red). Ubal is shown in foggy red. **b** Interaction surface with the ubiquitin globular domain of free YUH1 (light blue) and the complex (dark red). The arrows show the direction of the ubiquitin binding.

The binding surface with the ubiquitin globular domain is well converged in all ensemble structures of free YUH1 and similar to the complex, where two α-helices (α2 and α7) constitute the large groove (Fig. 4b). Meanwhile, the α7 helix in the free state structure is tilted slightly in the direction of widening the groove. Considering that the order parameters are relatively low in the end of this helix (Figure 1b), the free structure possesses a flexible wider groove than that of the complex for the sake of easy access of ubiquitin. Consequently, a conformational change presumably occurs when YUH1 cooperates with ubiquitin. The L5 loop, which was invisible in the YUH1-Uba1 crystal structure, is not well converged compared with other regions even in the free solution state (Fig. 3f).

### Specific interactions of the gating loop of YUH1 with ubiquitin

The NMR experiments and multi-state structure calculation elucidated that the gating loop and crossover loop possess large conformational variability. While the large dynamics of the crossover loop was speculated from previous reports on homologous proteins such as UCH-L3 ^18^, it has never been discussed that the gating loop passes transiently through and slips out of the crossover loop, resulting in the extensive conformational distribution of these regions. This result proposed a new hypothesis that its dynamics may correlate with an unknown function of the YUH1 gating loop. To elucidate the functional significance of this dynamics using a simpler model, we first performed an NMR titration experiment of a short peptide from the YUH1 N-terminus (1-10) [henceforth referred as YUH1(1-10)] with ubiquitin. It was expected that the titration of YUH1(1-10) with ubiquitin would reveal an interaction between the YUH1 gating loop and ubiquitin. Increasing the peptide ratio against ubiquitin, chemical shift perturbations of several peaks were explicitly observed (Fig. 5a and b). Surprisingly, the most significant chemical shift changes were observed not only for the C-terminus but also the center of the ubiquitin globular domain. In the crystal structure of the YUH1-Ubal complex the YUH1 gating loop is close and parallel to the ubiquitin C-terminus but rather far from the center of the ubiquitin globular domain (Supplementary Fig. 1). Large chemical shift differences cluster around the β-strands of ubiquitin, demonstrating that the gating loop of YUH1 specifically interacts with ubiquitin in a different orientation from the complex structure. The region showing large chemical shift perturbations is charged positively and forms a shallow groove on the surface of ubiquitin (Fig. 5c). Considering that the YUH1 gating loop is negatively charged, it may bind to this groove primarily by salt-bridge interactions. The dissociation constant calculated from the chemical shift perturbations was approximately 180 μM. Comparing this to the dissociation constant of the YUH1(C90S) mutant with ubiquitin that was reported as 43 nM ^34^, the affinity of YUH1(1-10) to ubiquitin is over 1000 times lower, suggesting that the interaction of the gating loop with ubiquitin might be transient.

**Fig. 5.**
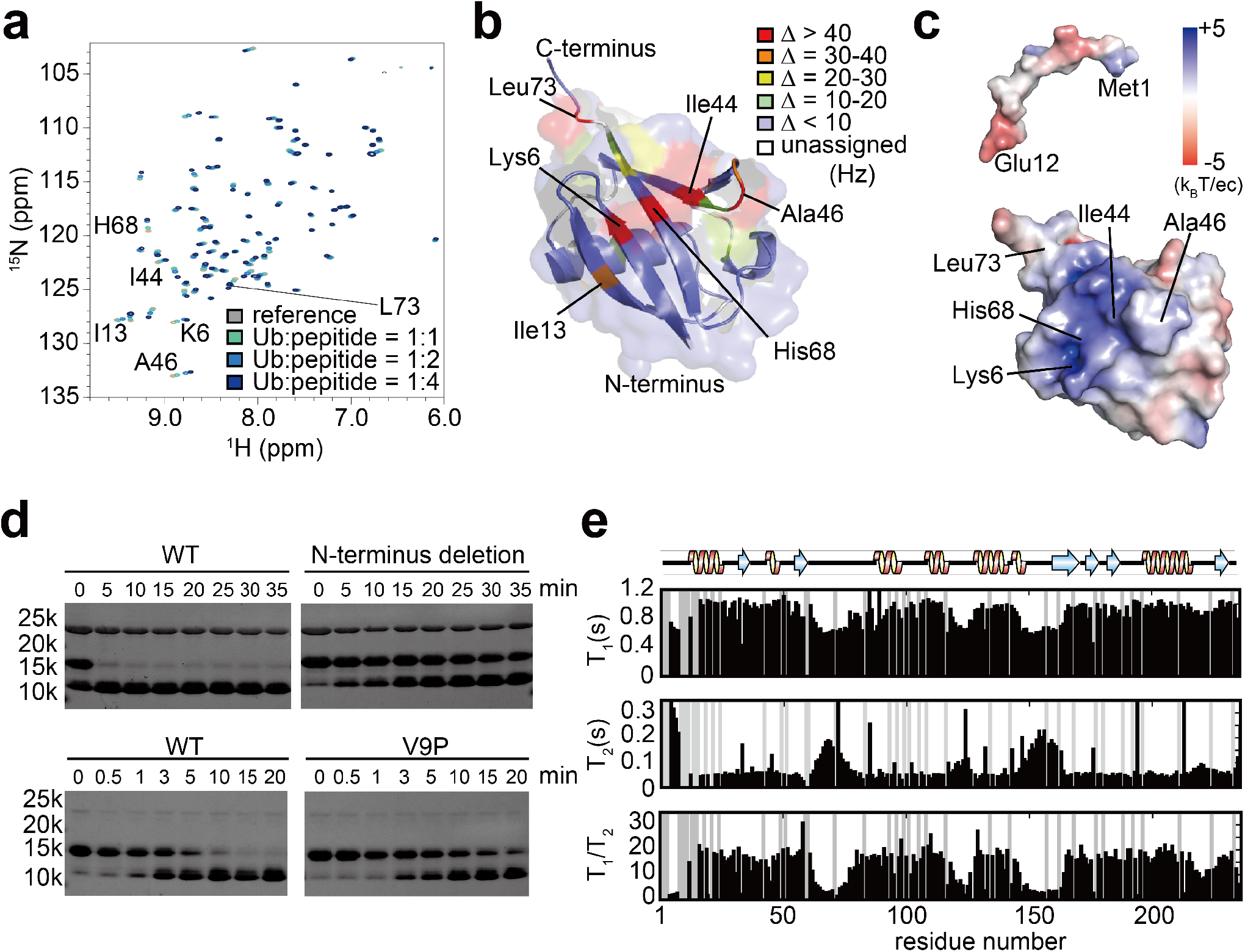
Functional assay of the YUH1 gating loop. **a** Overlays of 2D ^1^H-^15^N HSQC spectra from multipoint titrations of ^15^N-ubiquitin with the unlabeled N-terminus peptide of YUH1 (residues 1–10). The color codes of ^1^H-^15^N correlation cross-peaks at each titration point, showing the molar ratio of ubiquitin:peptide, are as follows: gray (1:0); green(1:1); light blue (1:2); blue (1:4). **b** Chemical shift changes *Δ* for ubiquitin upon binding of the N-terminus peptide, YUH1(1–10), mapped on a ribbon and surface model of the crystal structure of ubiquitin (PDB code:1UBQ). **c** Electrostatic potential for the N-terminal peptide and ubiquitin indicated by colored surface models. **d** YUH1 hydrolysis activity of Ub-Rec33 fusion protein. Reaction samples were collected at the given time points and subjected to SDS-PAGE electrophoresis for wild type (top and bottom left), the gating loop deletion mutant (top right) of YUH1, and the V9P mutton (bottom right). The reactions shown in the top and bottom panels were conducted at 25 and 20 °C, respectively. YUH1, Ub-Rec33 fusion protein, and cleaved Ub (in addition to a His_10_ tag at the Ub-Rec33 N-terminus) appear as a band at about 26, 15, and 11 kDa, respectively. Cleaved Rec33 is invisible on these electrophoresis gels because of its small size of 4 kDa. **e** *T*_1_, *T*_2_ relaxation times for backbone ^15^N resonances of YUH1. Residues without data are shown in grey.

### Hydrolysis activity of YUH1 and the role of the gating loop

The multi-state structure calculation showed that the gating loop possesses extraordinarily large dynamics but not an entirely random structure. To elucidate the correlation between the large-scale dynamics and ubiquitin-specific interaction of the gating loop with the YUH1 hydrolysis activity, we produced two mutants, an N-terminal deletion [henceforth referred as YUH1(11-236)] and the V9P mutant. The replacement of valine 9, located at the root of the N-terminal chain, to proline was expected to suppress the dynamics of the gating loop due to its rigidifying effect on the protein backbone. The *T*_1_/*T*_2_ relaxation for the V9P mutant (Fig. 5e) showed slightly longer *T*_2_ relaxation times for the gating loop than that for the wild type, indicating that the V9P mutant indeed influenced the gating loop dynamics. It is considered that the chemical exchange contribution to the *T*_2_ relaxation time is suppressed by the V9P mutant. On the basis of this physical property of the V9P mutant, the deubiquitination activities of both mutants as well as the wild type were evaluated at 25 °C by employing a ubiquitin fused with its C-terminus to the N-terminal 33 residues of *E. coli* RecA (Rec33) (henceforth referred to as Ub-Rec33). The SDS–PAGE analysis of the activity for the wild type YUH1 showed that the ubiquitin part was observed at the expected molecular size, showing that Ub-RecA33 was hydrolyzed precisely (Fig. 5d). In comparison with the wild type, YUH1(11-236) shows dramatically reduced hydrolysis activity. It did not completely cleave Ub-Rec33 even after 35 min while the wild type did it almost within 5 min, indicating that the gating loop significantly contributes to the enzymatic activity of YUH1. In contrast, under the same reaction conditions, the V9P mutant cleaved Ub-Rec33 as fast as the wild type (Supplementary Fig. 12). However, when repeating the experiments with a 20 times lower concentration of YUH1 and at a lower temperature of 20 °C, the V9P mutant failed to completely hydrolyze Ub-Rec33 even after 20 min, while the wild type achieved it within 15 min. This reduction of the enzymatic activity of YUH1 might be attributed to the change in the gating loop dynamics, which was demonstrated by the *T*_1_/*T*_2_ relaxation experiment. These biochemical analyses of YUH1(11–236) and the V9P mutant suggest that the gating loop and its dynamics are essential for the efficiency of the YUH1 function.

### Multiple conformations of the YUH1-ubiquitin complex

The multi-state structure determination and titration experiments of YUH1(1-10) with ubiquitin demonstrated that free YUH1 possesses multiple conformations and transiently interacts with different sites of ubiquitin. It indicates that the interaction of YUH1 with ubiquitin also includes multiple states. To investigate the binding states between YUH1 and ubiquitin, we recorded ^1^H-^15^N TROSY spectra for ^2^H, ^15^N-ubiquitin in the presence of non-labeled YUH1 at a low temperature slowing down molecular motions (283 K). Notably, two resonances were observed for K11, I13, T15, I36, D39, A46, K48, Q49, L67, H68, and V70 of ubiquitin (Fig. 6a), while only one resonance was observed for these residues in the spectra of ^2^H,^15^N-Ubal-unlabeled YUH1 complex. One of the split peaks in the ubiquitin-YUH1 complex was identical with that of the Ubal-YUH1 complex. These residues correspond to the interface of the YUH1-complex (Fig. 6b), suggesting that the interaction between ubiquitin and YUH1 is also composed of multiple-states in addition to the conformation observed in the Ubal-YUH complex.

**Fig. 6.**
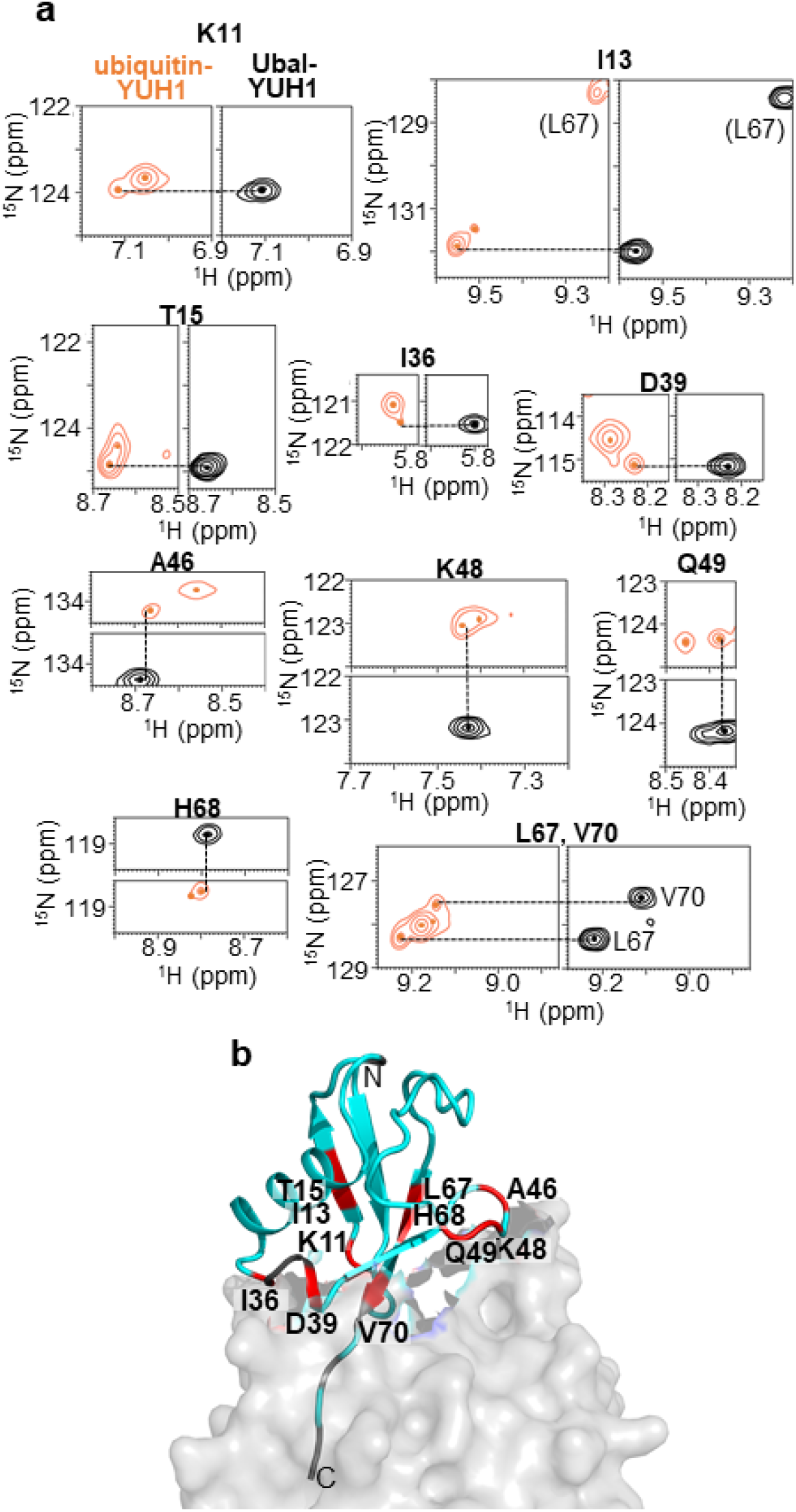
Multiple conformations of the YUH1-ubiquitin complex. **a** 2D ^1^H-^15^N TROSY spectrum of 0.85 mM ^2^H, ^15^N-ubiquitin with 1.45 mM unlabeled YUH1 (orange) and 0.5 mM [^2^H, ^15^N]Ubal-unlabeled YUH1 complex (black). Only the regions with K11, I13, T15, I36, D39, A46, K48, Q49, L67, H68, and V70 resonances are shown, and the center of the peaks are indicated with dots. Dashed lines represent the ^1^H or ^15^N chemical shifts of the resonances of Ubal-YUH. Full spectra are shown in Supplementary Figure 13. **b** Mapping of the residues (red) on Ubal that exhibited two resonances in the spectrum of the YUH1-Ubal complex (PDB ID: 1CMX). The residues with one resonance are color-coded in light blue, and the ones with no data, proline residues, or probably due to the line broadening, are colored in gray. YUH1 and Ubal are shown in the surface and ribbon diagrams, respectively.

## Discussion

The multi-state structure calculation with multiple NMR data sets presented here reveals large dynamics of the gating loop and the crossover loop of YUH1. In particular, the extensive motions of the gating loop first elucidated by this method allow us to find its potential functional significance. Based on the conformational ensemble, dynamics and interaction analysis by NMR, and biochemical evidence for the hydrolysis activity of the YUH1 mutants, we propose a new recognition mechanism of YUH1 with ubiquitin (Fig. 7). In the initiation of its reaction cycle, the free YUH1 exists in a largely fluctuating state for the regions surrounding the active site, the gating loop and crossover loop. The gating loop of YUH1 is predominantly in equilibrium state between two conformations, the closed form passing it through the crossover loop and the open form spreading widely by slipping it out of the loop. The titration experiment of ubiquitin with YUH1(1-10) demonstrates that the flexible gating loop specifically interacts with both the β-sheet in the globular domain and the C-terminus of ubiquitin. Truncation of the N-terminus in YUH1(11–236) dramatically reduces the enzymatic activity, and the V9P mutant shows moderately reduced activity. Longer *T*_*2*_ relaxation times for the gating loop in the V9P mutant indicate that a modulation of molecular motion for the gating loop occurs, which is presumably responsible for the reduced enzymatic activity. An earlier structural study of free UCH-L3 suggested that the function of its N-terminus is to prohibit non-specific enzymatic activity by covering the active center residue Cys90^18^. However, the reduction of enzymatic activity by modifications on the YUH1 gating loop indicates that it at least does not work for suppressing the non-specific catalytic cysteine nucleophile but rather enables its efficient reaction. Considering that the gating loop interacts not only with the ubiquitin C-terminus but also its globular domain, we propose a unique function of the gating loop that might capture a free ubiquitin and guide it to the binding and enzymatic active site. Based on the crystal structure of the YUH1-Ubal complex, the binding surface comprises roughly two regions, the one surrounding the enzymatic active site with the C-terminus of ubiquitin (site A) and the other interacting with the ubiquitin globular domain (site B). Since site B is composed of an open groove, its interaction is presumably more straightforward and faster than that of site A forming a narrow and deep hole. Thus, we assume that the initial step of the reaction is that the YUH1 gating loop captures the free ubiquitin and introduces it to site B. The ubiquitin-binding occurs as an induced fit where the site B groove narrows by the slight tilt of the α7 helix. During this process, the YUH1 gating loop would detach from the interaction with the β-sheet of ubiquitin, moves towards the active site and pass through the crossover loop. The ability for the gating loop to bind both with the ubiquitin C-terminus and the globular domain (Fig. 5c) indicates that the location shift of the YUH1 gating loop might occur together with the introduction of the ubiquitin C-terminus into the active site. Hence, the YUH1 gating loop might work as a guide of the ubiquitin C-terminus to the catalytic center. Considering that the temperature factors of the crystal structure of the YUH1-Ubal complex are relatively low in the active site, this area would be fixed after the passing process of the YUH1 gating loop and ubiquitin C-terminus through the crossover loop for the sake of an efficient and specific reaction. Finally, the enzyme performs the cleavage of the chemical adduct from the ubiquitin C-terminus. After the reaction, YUH1 releases ubiquitin and returns to the equilibrium between the open and closed states again. Observation of two sets of resonances for the YUH1-ubiquitin complex (Fig. 6) provides evidence for the above-described multi-step interaction, and it is tempting to speculate that the split resonances observed for the ubiquitin-YUH complex may correspond to the states before and after the passing process of the gating loop, or multiple binding sites of ubiquitin with the gating loop. Further NMR studies on the multiple conformations of the ubiquitin-YUH complex would be helpful for understanding the details of the ubiquitin-recognition process of YUH1.

**Fig. 7.**
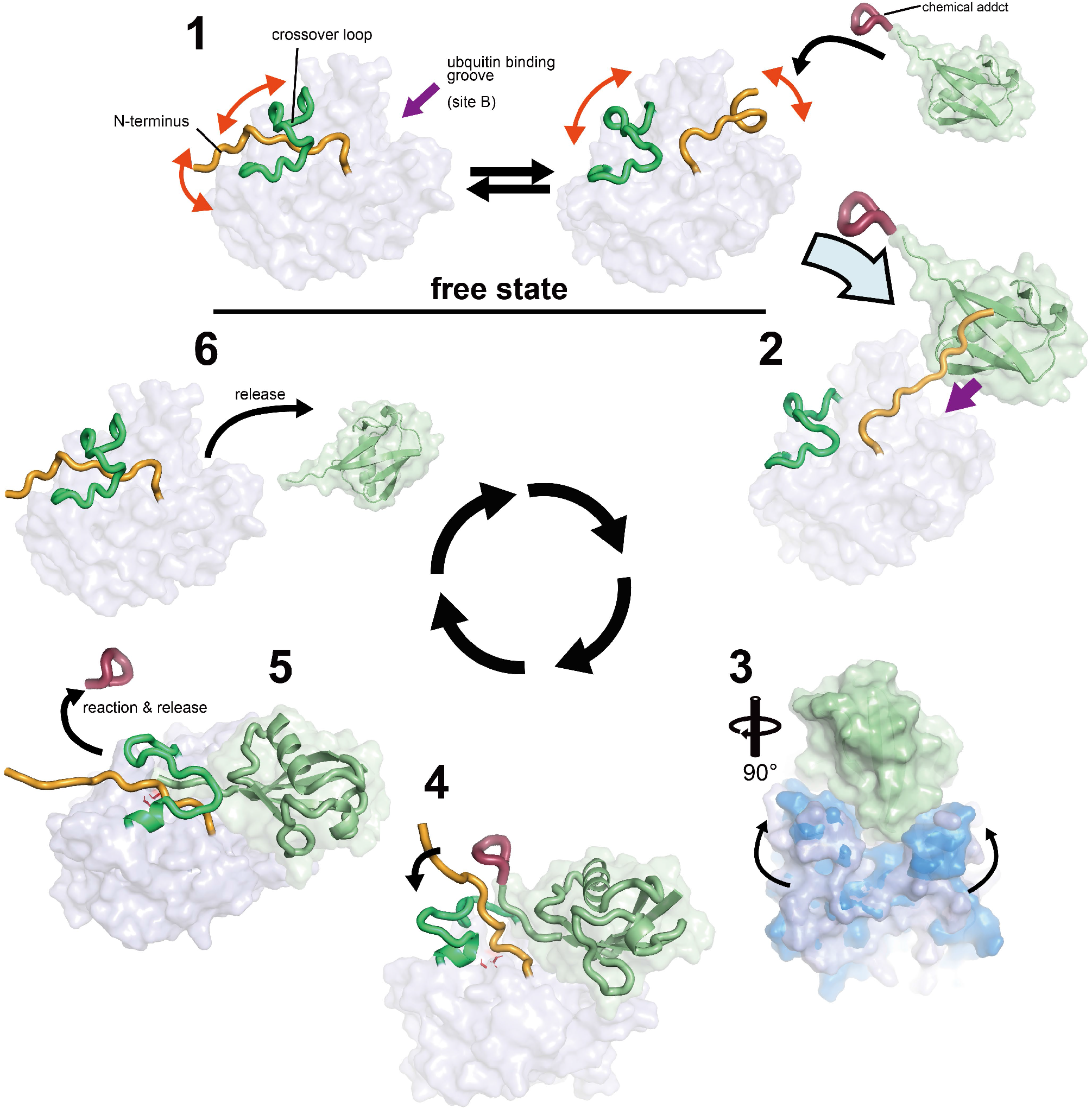
A proposed molecular recognition model for the ubiquitin C-terminal hydrolysis by YUH1. **1)** Free YUH1 is in the equilibrium state comprising both, the closed form with the gating loop passing through the crossover loop and the open form slipping the gating loop out of the loop. **2)** When ubiquitin is nearby, the YUH1 gating loop captures it as lariat by interacting with its globular domain. **3)** The ubiquitin globular domain binds at the site B of YUH1, and a conformational change primarily in helix α7 occurs. **4)** The gating loop dissociates from the ubiquitin globular domain and moves toward the enzymatic active center passing through the crossover loop along with the ubiquitin C-terminus. **5)** The crossover loop and gating loop of YUH1 anchor the ubiquitin C-terminus and hydrolyze a chemical adduct attached to the ubiquitin C-terminus. **6)** YUH1 releases ubiquitin and returns to the equilibrium state.

Using YUH1 as a model system, we demonstrated in this study that multi-state structure determination with multiple NMR data sets can be employed for general protein structural analysis and opens an avenue toward a comprehensive spatial description of 3D structures and the dynamics of biomacromolecules. Structure determination by X-ray crystallography may not allow to observe wide-spread conformational distributions such as the large motion of the YUH1 gating loop due to the suppression of dynamics in a crystal. It is crucial to observe the flexible regions in solution or under near-physiological conditions because of their susceptibility for their surrounding environment. Combining NMR CPMG relaxation dispersion and CEST with our method, one can also quantify the low population of each visualized conformation at various time-scales related to the dynamic behavior, which may not be possible even with cryo-electron microscopy. In this respect, NMR spectroscopy is better suited for such studies than the two other methods.

Our results demonstrate that, in addition to the conventional structural information such as NOEs and chemical shifts, the paramagnetic effects and RDCs are powerful tools to uncover the conformational distribution. Until recently, it was not trivial to collect accurate paramagnetic effects, in particular due to the large conformational flexibility of the metal-binding tags themselves. However, recent progress on lanthanoid-binding tags encourages us to observe the paramagnetic effects accurately and easily. Hence, we believe that it is now possible to perform ensemble structure determinations of various biomolecules and uncover molecular interactions and functions based on the comprehensive spatial description. Moreover, the current NMR methodologies allow us to measure biomolecules at atomic resolution even in living cells (in-cell NMR). Expanding our method to in-cell NMR can be a further useful tool to elucidate the physiological importance of protein dynamics through structural biology.

## Material and Methods

### Protein expression

The gene encoding *S. cerevisiae* YUH1 was constructed into the vector pET24a (Addgene) for over-expression in *Escherichia coli* as described previously^35^. Uniformly ^13^C,^15^N-labeled YUH1 protein was obtained by growing *E. coli* BL21 (DE3) strain harboring the expression vector at 37 °C in M9 minimal media, containing [^13^C_6_]-glucose (ISOTEC) and ^15^NH_4_Cl (ISOTEC) as the sole carbon and nitrogen sources, supplemented with 20 mM MgSO_4_, 0.1 mM CaCl_2_, 0.4 mg/ml thiamin, 20 μM FeCl_3_, salt mix (4 μM ZnSO_4_, 0.7 μM CuSO_4_, 1 μM MnSO_4_, 4.7 μM H_3_BO_3_), and 50 mg/l kanamycin. Protein expression was induced by adding 119 mg/l isopropyl β-D-thiogalactoside (IPTG) at an OD 600 nm of 0.6. After 6 h of further growth, cells were harvested and washed with a pH 7.5 lysis buffer (50 mM Tris-HCl, 2 mM EDTA, 5 mM DTT, and 100 mM NaCl). Uniformly ^15^N-labeled YUH1 was produced identically except using [^12^C_6_]-glucose. Uniformly single ^15^N-labeled and double ^13^C,^15^N-labeled YUH1 proteins with 50% random fractional deuteration (50% ^2^H/^15^N-YUH1) were obtained by growing cells in M9 medium containing 60% ^2^H_2_O/40% ^1^H_2_O. Ile/Leu/Val-methyl-selectively ^1^H/^13^C-labeled and Phe/Tyr/Trp-aromatic ring-selectively ^1^H-labeled YUH1 with perdeuterated background (henceforth referred as ILVFYW-YUH1) was obtained by combining the methods described by Rosen *et al*. ^36^ and Rajesh *et al*. ^37^. The ILVFYW-YUH1 sample was obtained growing the *E. coli* AB2826(DE3)/shiA strain harboring the expression vector in 100% D_2_O M9 medium containing 2 g/L [^13^C_6_, C-^2^H_7_] glucose (ISOTEC), 1 g/L ^15^NH_4_Cl, 100 mg/L [u-^13^C, 3,3-^2^H] α-ketobutyrate (CIL), 100 mg/L [u-^13^C, 3-_2_H] α-ketoisovalerate (CIL), to which 240 mg/L unlabeled shikimic acid was added prior to protein induction.

Ub-Rec33 fusion protein in addition to a His_10_ tag at the gating loop was constructed into the expression vector pET14b and over-expressed in the *E. coli* BL21 (DE3) strain. The human ubiquitin gene was inserted into a pET24a vector for overexpression in the *E. coli* BL21 Star (DE3) strain. Non-labelled samples of Ub-Rec33 fusion and human ubiquitin were prepared by growing the bacteria at 37 °C in LB media. Protein expression was induced in the same way as for wild type YUH1.

^2^H,^15^N-yeast ubiquitin was expressed as previously described^38^. The short peptide from the YUH1 gating loop (1-10) was commercially obtained from Toray Research Center Incorporated.

### Protein purification

All the procedures described below were carried out at 4 °C unless stated otherwise. All isotope-labelled YUH1 samples as well as the mutants were purified in the same way. The cells dispersed in the lysis buffer were disrupted by sonication for 30 min on ice with hen egg lysozyme (0.1 mg/ml). The cell debris was clarified by centrifugation at 14,000 *g* for 1 h. The supernatant was loaded onto a 25 ml DEAE-Sepharose Fast Flow (Cytiva) anion exchange equilibrated with buffer A [50 mM Tris-HCl (pH 7.5), 2 mM EDTA, 5 mM dithiothreitol (DTT)]. After washing the column with buffer A until sufficiently low UV absorption at 280 nm, the YUH1 protein was eluted by linearly increasing the concentration of NaCl from 0 to 1 M with a flow rate of 3 ml/min in buffer A. The fractions containing the target protein were concentrated to 5 ml with Amicon Ultra-15 10 kDa (Merck). The concentrated sample was loaded onto a 320 ml HiLoad Superdex 75 (GE Healthcare Life Science) gel filtration column with a flow rate of 1 ml/min using FPLC systems (AKTA pure 25 and AKTA FPLC, GE Healthcare Life Science). The 5 ml sample concentrated, from the fractions involving the target proteins, with Amicon Ultra-15 10 kDa was loaded on a Resource Q (GE Healthcare Life Science) anion-exchange column equilibrated with buffer A using the FPLC systems. After washing the column with 30 ml of buffer A, the YUH1 protein was eluted by linearly increasing the concentration of NaCl from 150 mM to 1 M with a flow rate of 0.5 ml/min in buffer A. The purity of the YUH1 samples was confirmed in each step by SDS-PAGE. Protein concentrations were determined by NanoDrop 2000 (ThermoFisher) measuring UV absorption at 280 nm. Protein samples for NMR measurements were concentrated and dissolved in NMR buffer (90% ^1^H_2_O/ 10% ^2^H_2_O containing 50 mM Na_2_HPO_4_-NaH_2_PO_4_, pH 6.0, 100 mM NaCl, 10 mM DTT). Yeast ubiquitin and Ubal-YUH1 complex was prepared as described previously^25,35^.

For purification of human ubiquitin, the harvested cells were re-suspended in the lysis buffer [50 mM Tris-HCl (pH 7.4), 1 mM DTT and 1 mM EDTA] and lysed by sonication. The cleared lysate was collected after centrifugation, and then heated to 80 °C for 15 minutes. Precipitated proteins were removed by centrifugation. The supernatant was loaded onto a DEAE-Sepharose FF column (GE Healthcare Life Science) pre-equilibrated with 50 mM acetate buffer. The column was washed with 5 column volumes of the same buffer, followed by elution with a linear gradient of 0-1 M NaCl. The fractions containing ubiquitin were concentrated by Amicon Ultra-15 3 kDa (Merck) and then loaded onto a HiLoad 16/60 Superdex 75 column (GE Healthcare Life Science) pre-equilibrated with the preparation buffer [20 mM Tris-HCl (pH 7.4), 1 mM DTT and 1 mM EDTA]. The final ubiquitin fractions were dissolved in a buffer [20 mM MES (pH 6.2), TCEP 0.3 mM, NaCl 100 mM].

For purification of Ub-Rec33 fusion protein, the cells that over-expressed the fusion proteins were sonicated by the above procedure in 50 mM Tris buffer [50 mM Tris-HCl (pH 7.4), 1mM DTT, 1mM EDTA]. The supernatant was loaded onto a 5 ml Ni-NTA agarose bead slurry (Qiagen) equilibrated with equilibration buffer [20 mM Tris-HCl (pH 8.0), 300 mM NaCl, 20 mM imidazole]. After sufficient washing of the column with equilibration buffer, the fusion protein was eluted by linearly increasing the concentration of imidazole from 20 to 500 mM with a flow rate of 1 ml/min in equilibration buffer. The eluted sample was exchanged into 20 mM Tris buffer and concentrated to 5 ml by Amicon Ultra-15 3 kDa (Merck). The sample was purified by 320 ml of HiLoad Superdex with a flow rate of 1 ml/min using the FPLC systems.

### Preparation of YUH1 mutants

For the measurements of paramagnetic effects, we introduced C90S and an additional cysteine mutant, N5C, S104C, N140C, and N225C. The DNA fragments encoding the YUH1 genes containing these mutants were synthesized, respectively. Generally, while the PRE and PCS effects reach maximally 30-40 Å, NMR signals nearby the paramagnetic centers disappear due to strong paramagnetic relaxation. Thus, in order to clearly observe the paramagnetic effects surrounding the crossover loop and gating loop, we designed cysteine mutants on residues that are approximately 10–15 Å away from the active site based on the crystal structure of the YUH1-Ubal complex. The DNA fragments were inserted into a pET24a vector for overexpression in the *E. coli* BL21 (DE3) strain. Uniformly ^15^N-labelled samples were prepared by growing the bacteria in M9 medium containing ^15^NH_4_Cl. These samples were purified in the same way as the wild type YUH1.

### Attachment of paramagnetic tags to ^15^N-labelled YUH1 mutant proteins

The spin-labeled MTSL samples of the cysteine variants were obtained by mixing with a five-fold excess of MTSL (Toronto Research Chemicals, Ontario, Canada) in a phosphate buffer [50 mM sodium phosphate, 100 mM NaCl at pH 8.0] and incubating for approximately 15 h at 25 °C. The protein solution was filtered and concentrated using Amicon Ultra-15 10kDa.

The [Mn^2+^(maleimide-DOTA)] samples were obtained by reacting with a three-fold excess of maleimide-DOTA (CheMatech) in a phosphate buffer [20 mM sodium phosphate, 0.5 mM DTT at pH 6.5] and a two-fold excess of MnCl_2_ for approximately 45 h at 35 °C. Subsequently, the protein samples were incubated with a two-fold excess of paramagnetic metal ions, and exchanged to the NMR buffer in the same way.

For the 4PS-PyMTA-attached YUH1 samples, a five-fold excess of 4PS-PyMTA (Taiyo Nippon Sanso Corp.) was added to the protein solution in a Tris buffer [20 mM Tris-HCl (pH 8.0), 0.3 mM TCEP], and incubated for 3 days at 30 °C. Equal amounts of paramagnetic (Cu^2+^, Mn^2+^) or diamagnetic (Zn^2+^) metal ions were then added to the protein solutions. The tagging reactions for 4PS-PyMTA failed at positions 104, 140, and 225 as manifested by non-observation of chemical shift or peak intensity changes at peaks corresponding to these residues.

The DOTA-M8-SPy-Ln-attached samples were prepared by adding a three-fold excess of DOTA-M8-SPy containing paramagnetic (Dy^3+^) or diamagnetic (Lu^3+^) metal ions (Taiyo Nippon Sanso Corp.) and incubating in a buffer [10 mM sodium phosphate (pH 5.0)] for a day at 25 °C. The DO3MA-3BrPy-Ln-tagged samples were obtained by incubating with a five-fold excess of DO3MA-3BrPy containing paramagnetic metal ions, Tm^3+^, Tb^3+^, Lu^3+^, and Yb^3+^ (Taiyo Nippon Sanso Corp.) in a buffer [20 mM Tris (pH 8.0), 0.2 mM TCEP] for a day at 25 °C. The DOTA-M8-SPy tag with diamagnetic and paramagnetic metals was conjugated through a disulfide bond to the YUH1 cysteine mutants. The DO3MA-3BrPy-attached sample was obtained by connecting through a thioether bond. In paramagnetically shifted spectra, resolved peaks were assigned by visual inspection considering the uniform direction of peak shifts and, if necessary, the expected size of peak shifts predicted from the Δχ tensor estimated from the already unambiguously assigned peaks.

All final protein solutions for PRE and PCS were adjusted to a concentration of 0.3 mM in the NMR buffer.

### YUH1 hydrolysis of ubiquitin fusion

Hydrolysis activity of YUH1 and YUH1 mutants was tested at 20 and 25 °C using the engineered ubiquitin with Rec33 as a substrate. The reactions were performed at different time points in a Tris buffer [Tris-HCl (pH 8.5), 0.5 mM EDTA, 1 mM DTT]. The reaction activity was analyzed by 13.5 % Tricine-SDS-PAGE.

### NMR spectroscopy

NMR experiments were performed at 30°C probe temperature on Bruker DRX, AVANCE, and AVANCE-III 600 and 700 MHz spectrometers equipped with pulsed field gradient triple-resonance cryoprobes. Spectra were processed with the Azara software package^39^. For the 3D data, the two-dimensional maximum entropy method (2D MEM) or Quantitative Maximum Entropy (QME) ^40^ was applied to obtain resolution enhancement for the indirect dimensions. NMR spectra were visualized and analyzed using the CcpNmr Analysis 2.5.0 software^41^. Backbone and side-chain resonance assignments were achieved by analyzing 3D triple-resonance NMR spectra, HNCA, HN(CO)CA, CBCANH, CBCA(CO)NH. HBHA(CBCACO)NH, H(CCCO)NH, CC(CO)NH, and HCCH-TOCSY, measured on 50%-^2^H/^13^C/^15^N-YUH1 samples. The ^1^H/^13^C resonance assignment of isoleucine δ1, leucine δ and valine γ methyl groups were confirmed unambiguously by analyzing 3D CC(CO)NH, CCNH and H(CCCO)NH spectra measured on the ILVFYW-YUH1 sample. The ^1^H resonances of phenylalanine, tyrosine and tryptophan aromatic rings were assigned by analyzing 2D ^1^H-^1^H TOCSY spectra measured on ILVFYW-YUH1 samples with reference to intraresidual and sequential NOEs. For the collection of NOE-derived distance restraints, 3D ^15^N-separated and 3D ^13^C-separated NOESY-HSQC spectra were measured on uniformly ^15^N-labeled or ^13^C-labeled samples, respectively. In addition, 3D ^1^H^aromatic^-edited/^15^N-separated NOESY-HSQC, 3D ^13^C-separated NOESY-HSQC, 3D ^13^C/^15^N-separated HSQC-NOESY-HSQC, and 3D ^13^C/^13^C-separated HSQC-NOESY-HSQC spectra were measured on the ILVFYW-YUH1 sample. A 100 ms NOE mixing period was employed for the 3D NOESY experiments. For restraints for hydrogen bonds, we measured scalar-couplings (^3h^*J*_N-C’_) recording a long-range TROSY-HNCO experiment on the ILVFYW sample. ^1^H-^15^N TROSY spectra were recorded for 450 mL of 1 mM ^2^H,^15^N-yeast ubiquitin supplemented with 80 μL of 9.6 mM unlabeled YUH1 and for 0.5 mM ^2^H,^15^N-Ubal-unlabeled YUH1 complex, in a buffer containing 50 mM NaPi, pH6.0, 100 mM NaCl, 1 mM DSS, and 10 % D_2_O at 283 K with Bruker Avance 600 spectrometers.

The longitudinal relaxation times (*T*_1_), the transverse relaxation times (*T*_2_), and the steady-state heteronuclear {^1^H}-^15^N NOEs were measured at 30 °C using uniformly ^15^N-labeled YUH1 on a Bruker Avance III 600 MHz spectrometer equipped with a cryogenic H/C/N triple-resonance probehead. Each experiment was acquired in a pseudo 3D manner. 8 relaxation delays in the range 15–1005 ms and 6 delays between 14–110 ms were used for the *T*_1_ and *T*_2_ experiments, respectively. NOE ratios were obtained from intensities in experiments recorded with (2 s relaxation delay followed by 3 s saturation) and without (relaxation delay of 5 s) saturation. The overall rotational correlation time, effective correlation times, and order parameters were obtained by the program relax 4.0.3^42^ with spectral density functions as defined by the Lipari-Szabo model-free approach. A theoretical rotational correlation time (*τ*_*c*_) was calculated by the Stokes-Einstein-Debye relation,

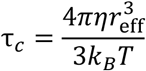

where *η* is the viscosity, *r*_eff_ is the effective hydrodynamic radius of a molecule, *k*_B_ is the Boltzmann constant, and *T* is the temperature. The rotational correlation time of YUH1 was estimated with *T* = 298.0 K, *η* = 0.890 mPa, and *r*_eff_ = 27.1 Å.

For RDC data collection, alignment of the protein was achieved using 20.0 mg/ml Pf1 phage (ASLA Biotech) in the NMR buffer. Measurements of ^1^*J*_HN_ splittings were carried out with 2D ^15^N-^1^H IPAP HSQC for the isotropic and aligned samples.

The titration experiments of ubiquitin with the N-terminus (1-10) of YUH1 were performed in a buffer [20 mM MES (pH 6.2), TCEP 0.3 mM, NaCl 100 mM] at 25 °C. ^15^N-ubiquitin was successively titrated with unlabeled N-terminus of YUH1 in molar ratios of 1:0.5, 1:1, 1:2, 1:4, 1:8, 1:16, and 1:32, respectively. 2D ^1^H-_15_N HSQC spectra were measured at each titration point. Chemical shift differences, *Δ*, were analyzed by *Δ* = (*Δ* _H2_ + *Δ*N_2_) ^1/2^, where *Δ* _H_ and *Δ* _N_ are the differences in Hz for the backbone amide ^1^H and ^15^N chemical shifts between the two conditions. 1 ppm corresponds to 600.13 Hz for ^1^H and 60.81 Hz for ^15^N. The dissociation constants for each residue were calculated by non-linear regression analysis with the equation^43^:

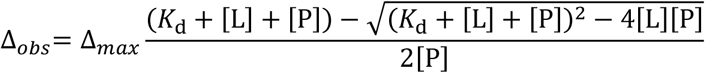

where *Δ*_obs_ is the observed chemical shift perturbation, *Δ*_max_ is the maximum chemical shift perturbation, *K*_d_ is the dissociation constant, and [L] and [P] are the ligand and protein concentrations respectively. The non-linear least squares fitting was performed by a homemade python script with the Levenberg-Marquardt algorithm. The assignment of ubiquitin signals was performed based on assignments deposited in the Biological Magnetic Resonance Data Bank (BMRB) (entry number:27356)

### Conventional single-state structure calculation

We manually analyzed intra-residual and long-range NOEs involving methyl and aromatic protons on 3D ^15^N- and ^13^C-separated NOESY-HSQC, and ^13^C/^13^C-separated HMQC-NOESY-HMQC spectra. Based on the manual NOE assignments, additional analysis was automatically performed by the program CYANA with the use of automated NOE assignment and torsion angle dynamics^14^.

The peak position tolerance was set to 0.03 ppm for the ^1^H dimension and to 0.3 ppm for the ^13^C and ^15^N dimensions. Backbone torsion angle restraints were obtained from chemical shifts with the program TALOS-N^44^. To obtain distance restraints, the PRE experiments were processed similar to the protocol of Battiste and Wagner^45^. On the basis of the peak intensities *I*_para_ and *I*_dia_ in the paramagnetic and diamagnetic state, respectively, three classes of distance restraints from PRE data were prepared for structure calculations as described^45^, except that a tolerance of 6 Å instead of 4 Å was applied and a lower bound of 6 Å was used if *I*_para_/*I*_dia_ > 0.85. The components of the magnetic susceptibility tensor and the metal ion position were iteratively determined by fitting experimental PCS to PCS backcalculated from intermediate structures in the calculation. The alignment tensor for RDCs was also determined fitting the experimental data to the primary YUH1 structure. The PCS and RDC tensor parameter fittings were performed with the Levenberg–Marquardt algorithm and singular value decomposition, respectively.

Seven cycles of combined automated NOE assignment and structure calculation by simulated annealing in torsion angle space and a final structure calculation using only unambiguously assigned distance restraints were performed. Calculations were started from 100 conformers with random torsion angle values, simulated annealing with 50,000 torsion angle dynamics steps was applied, and the 20 conformers with the lowest final target-function values were analyzed.

### Multi-state structure calculation

For the ensemble-averaged calculations, two to eight structural states of the entire protein were calculated simultaneously, excluding steric repulsion between atoms of different states. NOE distance restraints derived from the conventional automated NOE assignment described above were applied as 1/*r*^6^ averages of the corresponding distances in the individual states. Torsion angle restraints, RDCs, PREs, and PCSs were applied by averaging the corresponding quantities in the individual states. The multi-state structure calculation protocol used multiple copies of the amino acid sequence of YUH1 connected with invisible linkers and performed simulated annealing by torsion angle dynamics with CYANA to acquire ensemble structures that fulfill experimental restraints. We employed weak artificial harmonic restrains between corresponding atoms of the individual chains in order to keep states together unless multiple different states are required to fulfill the experimental restraints in regions with large dynamic. To maintain the conformers of the individual states into identical positions as far as permitted by the experimental restraints, weak upper distance bounds of 1.2 Å were employed on all distances between the same nitrogen and carbon atoms in different states. Calculations started from 2500 random conformers with five states, and the 500 conformers with the lowest final target-function values were selected for analysis. By dividing each multi-state conformer into five individual structures, a total 2500 states were obtained as the final ensemble structures of YUH1 and analyzed by PCA. The fractions of variance explained by the first four principal components were 0.649, 0.162, 0.087, and 0.037. As the cumulative fraction of variance for the first and second components reached above 0.8, we carried out k-means clustering only for the first and second principal component assuming five clusters. One conformer located at the cluster center and three randomly selected ones were obtained from each cluster to form the 20 conformers that were employed as the final representative structures. For PDB deposition, the 20 conformers were subjected to restrained energy minimization in explicit solvent against the AMBER force field^46,47^, using the program OPALp ^48^.

### Cross-validation test

We randomly divided the full experimental data into ten subsets of equal size, in which a subset was used for validation, and the rest was for a test calculation^49^. The number of the models was evaluated by the average of ten independent tests repeated with different combinations. Here, we performed multi-state structure calculations ranging from two to eight conformational states.

## Data availability

Atomic coordinates of free YUH1 have been deposited in the Protein Data Bank with the accession code 7EN4. All NMR chemical shifts have been deposited in the BioMagResBank with accession numbers 36420. All other data are available from the corresponding author upon reasonable request.

## Acknowledgment

The authors thank Dr. Tsutomu Terauchi (Taiyo Nippon Sanso Corp.) and Dr. Youhei Kawabata for useful discussions on paramagnetic tags and NMR relaxation, respectively. We gratefully acknowledge financial support from the Funding Program for Core Research for Evolutional Science and Technology (CREST; JPMJCR13M3) from the Japan Science and Technology Agency (JST), Grants-in-Aid for Scientific Research (JP15K06979 to T.I., JP17K07312 to P.G., JP19H05645 to Y.I.) and Scientific Research on Innovative Areas (JP26102538, JP25120003, JP16H00779 and JP21K06114 to T.I., JP15H01645, JP16H00847,JP17H05887 and JP19H05773 to Y.I.) from the Japan Society for the Promotion of Science (JSPS), Shimazu foundation, and the Precise Measurement Technology Promotion Foundation. NMR experiments with a Bruker Avance III 700 MHz spectrometer were performed at the RIKEN NMR Platform supported by the Ministry of Education, Culture, Sports, Science and Technology (MEXT) of Japan.

## Author Contributions

P.G., Y.I and T.I. designed the research and wrote the manuscript. M.O., Y.T., E.N., S.R. and T.I. conducted the research including sample preparation, data acquisition, resonance assignment and structure calculation. T.M. prepared mutants of YUH1. H.Y. and T. Kigawa helped with paramagnetic NMR measurements. T.U. and S.I. designed the research for the YUH1-ubiquitin interaction. H.O. and T.U. conducted sample preparations and NMR measurements for the YUH1-ubiquitin complex. T. Kohno prepared wild type YUH1.

## Notes

### Competing Interest Statement

The authors have declared no competing interest.

## References

1. Goh, C. S., Milburn, D. & Gerstein, M. Conformational changes associated with protein-protein interactions. Curr. Opin. Struct. Biol. 14, 104–109 (2004).

2. Wright, P. E. & Dyson, H. J. Intrinsically disordered proteins in cellular signalling and regulation. Nat. Rev. Mol. Cell Biol. 16, 18–29 (2015).

3. Ikeya, T., Güntert, P. & Ito, Y. Protein Structure Determination in Living Cells. Int. J. Mol. Sci. 20 (2019).

4. Ito, Y.D.V.; Shirakawa, M. In-cell NMR Spectroscopy: From Molecular Sciences to Cell Biology. (The Royal Society of Chemistry, 2019).

5. Korzhnev, D. M. & Kay, L. E. Probing invisible, low-populated States of protein molecules by relaxation dispersion NMR spectroscopy: an application to protein folding. Acc. Chem. Res. 41, 442–451 (2008).

6. Fawzi, N. L., Ying, J., Ghirlando, R., Torchia, D. A. & Clore, G. M. Atomic-resolution dynamics on the surface of amyloid-beta protofibrils probed by solution NMR. Nature 480, 268–272 (2011).

7. Vallurupalli, P., Bouvignies, G. & Kay, L. E. Studying “invisible” excited protein states in slow exchange with a major state conformation. J. Am. Chem. Soc. 134, 8148–8161 (2012).

8. Nodet, G. et al. Quantitative description of backbone conformational sampling of unfolded proteins at amino acid resolution from NMR residual dipolar couplings. J. Am. Chem. Soc. 131, 17908–17918 (2009).

9. Tang, C., Schwieters, C. D. & Clore, G. M. Open-to-closed transition in apo maltose-binding protein observed by paramagnetic NMR. Nature 449, 1078–1082 (2007).

10. Liu, W., Liu, X., Zhu, G., Lu, L. & Yang, D. A Method for Determining Structure Ensemble of Large Disordered Protein: Application to a Mechanosensing Protein. J. Am. Chem. Soc. 140, 11276–11285 (2018).

11. Kooshapur, H., Schwieters, C. D. & Tjandra, N. Conformational Ensemble of Disordered Proteins Probed by Solvent Paramagnetic Relaxation Enhancement (sPRE). Angew. Chem. Int. Ed. Engl. 57, 13519–13522 (2018).

12. Vögeli, B., Kazemi, S., Güntert, P. & Riek, R. Spatial elucidation of motion in proteins by ensemble-based structure calculation using exact NOEs. Nat. Struct. Mol. Biol. 19, 1053–1057 (2012).

13. Clore, G. M. & Iwahara, J. Theory, practice, and applications of paramagnetic relaxation enhancement for the characterization of transient low-population states of biological macromolecules and their complexes. Chem. Rev. 109, 4108–4139 (2009).

14. Güntert, P. & Buchner, L. Combined automated NOE assignment and structure calculation with CYANA. J. Biomol. NMR 62, 453–471 (2015).

15. Liu, C. C., Miller, H. I., Kohr, W. J. & Silber, J. I. Purification of a Ubiquitin Protein Peptidase from Yeast with Efficient Invitro Assays. J. Biol. Chem. 264, 20331–20338 (1989).

16. Song, L. & Luo, Z. Q. Post-translational regulation of ubiquitin signaling. J. Cell Biol. 218, 1776–1786 (2019).

17. Nijman, S. M. et al. A genomic and functional inventory of deubiquitinating enzymes. Cell 123, 773–786 (2005).

18. Johnston, S. C., Riddle, S. M., Cohen, R. E. & Hill, C. P. Structural basis for the specificity of ubiquitin C-terminal hydrolases. EMBO J. 18, 3877–3887 (1999).

19. Das, C. et al. Structural basis for conformational plasticity of the Parkinson’s disease-associated ubiquitin hydrolase UCH-L1. Proc. Natl. Acad. Sci. U S A 103, 4675–4680 (2006).

20. Misaghi, S. et al. Structure of the ubiquitin hydrolase UCH-L3 complexed with a suicide substrate. J. Biol. Chem. 280, 1512–1520 (2005).

21. Nishio, K. et al. Crystal structure of the de-ubiquitinating enzyme UCH37 (human UCH-L5) catalytic domain. Biochem. Biophys. Res. Commun. 390, 855–860 (2009).

22. Artavanis-Tsakonas, K. et al. Characterization and structural studies of the Plasmodium falciparum ubiquitin and Nedd8 hydrolase UCHL3. J. Biol. Chem. 285, 6857–6866 (2010).

23. Navarro, M. F., Carmody, L., Romo-Fewell, O., Lokensgard, M. E. & Love, J. J. Characterizing substrate selectivity of ubiquitin C-terminal hydrolase-L3 using engineered alpha-linked ubiquitin substrates. Biochemistry 53, 8031–8042 (2014).

24. Johnston, S. C., Larsen, C. N., Cook, W. J., Wilkinson, K. D. & Hill, C. P. Crystal structure of a deubiquitinating enzyme (human UCH-L3) at 1.8 Å resolution. EMBO J. 16, 3787–3796 (1997).

25. Rajesh, S. et al. Ubiquitin binding interface mapping on yeast ubiquitin hydrolase by NMR chemical shift perturbation. Biochemistry 38, 9242–9253 (1999).

26. Cordier, F. & Grzesiek, S. Direct Observation of Hydrogen Bonds in Proteins by Interresidue 3hJNC’ Scalar Couplings. Journal of the American Chemical Society 121, 1601–1602 (1999).

27. Popp, M. W., Artavanis-Tsakonas, K. & Ploegh, H. L. Substrate filtering by the active site crossover loop in UCHL3 revealed by sortagging and gain-of-function mutations. J Biol Chem 284, 3593–3602 (2009).

28. Häussinger, D., Huang, J. R. & Grzesiek, S. DOTA-M8: An extremely rigid, high-affinity lanthanide chelating tag for PCS NMR spectroscopy. J. Am. Chem. Soc. 131, 14761–14767 (2009).

29. Yang, Y. et al. A Reactive, Rigid Gd(III) Labeling Tag for In-Cell EPR Distance Measurements in Proteins. Angew. Chem. Int. Ed. Engl. 56, 2914–2918 (2017).

30. Yang, Y., Wang, J. T., Pei, Y. Y. & Su, X. C. Site-specific tagging proteins via a rigid, stable and short thiolether tether for paramagnetic spectroscopic analysis. Chem. Commun. 51, 2824–2827 (2015).

31. Theillet, F. X. et al. Structural disorder of monomeric alpha-synuclein persists in mammalian cells. Nature 530, 45–50 (2016).

32. Berliner, L. J., Grunwald, J., Hankovszky, H. O. & Hideg, K. A novel reversible thiol-specific spin label: Papain active site labeling and inhibition. Anal. Biochem. 119, 450–455 (1982).

33. Ottiger, M., Delaglio, F. & Bax, A. Measurement of J and dipolar couplings from simplified two-dimensional NMR spectra. J. Magn. Reson. 131, 373–378 (1998).

34. Igarashi, S., Osawa, M., Takeuchi, K., Ozawa, S. & Shimada, I. Amino acid selective cross-saturation method for identification of proximal residue pairs in a protein-protein complex. J. Am. Chem. Soc. 130, 12168–12176 (2008).

35. Sakamoto, T. et al. An NMR analysis of ubiquitin recognition by yeast ubiquitin hydrolase: evidence for novel substrate recognition by a cysteine protease. Biochemistry 38, 11634–11642 (1999).

36. Rosen, M. K. et al. Selective methyl group protonation of perdeuterated proteins. J. Mol. Biol. 263, 627–636 (1996).

37. Rajesh, S. et al. A novel method for the biosynthesis of deuterated proteins with selective protonation at the aromatic rings of Phe, Tyr and Trp. J. Biomol. NMR 27, 81–86 (2003).

38. Matsumoto, M., Ueda, T. & Shimada, I. Theoretical analyses of the transferred cross-saturation method. J Magn Reson 205, 114–124 (2010).

39. Boucher, W. Azara, v2.7 http://www.bio.cam.ac.uk/azara/.

40. Hamatsu, J. et al. High-resolution heteronuclear multidimensional NMR of proteins in living insect cells using a baculovirus protein expression system. J. Am. Chem. Soc. 135, 1688–1691 (2013).

41. Vranken, W. F. et al. The CCPN data model for NMR spectroscopy: development of a software pipeline. Proteins 59, 687–696 (2005).

42. Bieri, M., d’Auvergne, E. J. & Gooley, P. R. relaxGUI: a new software for fast and simple NMR relaxation data analysis and calculation of ps-ns and mus motion of proteins. J. Biomol. NMR 50, 147–155 (2011).

43. Morton, C. J. et al. Solution structure and peptide binding of the SH3 domain from human Fyn. Structure 4, 705–714 (1996).

44. Shen, Y. & Bax, A. Protein backbone and sidechain torsion angles predicted from NMR chemical shifts using artificial neural networks. J. Biomol. NMR 56, 227–241 (2013).

45. Battiste, J. L. & Wagner, G. Utilization of site-directed spin labeling and high-resolution heteronuclear nuclear magnetic resonance for global fold determination of large proteins with limited nuclear overhauser effect data. Biochemistry 39, 5355–5365 (2000).

46. Cornell, W. D. et al. A second generation force field for the simulation of proteins, nucleic acids, and organic molecules. J. Am. Chem. Soc. 117, 5179–5197 (1995).

47. Ponder, J. W. & Case, D. A. Force Fields for Protein Simulations. Adv. Protein Chem. 66, 27–85 (2003).

48. Koradi, R., Billeter, M. & Güntert, P. Point-centered domain decomposition for parallel molecular dynamics simulation. Comput. Phys. Commun. 124, 139–147 (2000).

49. Vögeli, B., Olsson, S., Riek, R. & Güntert, P. Complementarity and congruence between exact NOEs and traditional NMR probes for spatial decoding of protein dynamics. J. Struct. Biol. 191, 306–317 (2015).

50. Gottstein, D., Kirchner, D. K. & Güntert, P. Simultaneous single-structure and bundle representation of protein NMR structures in torsion angle space. J. Biomol. NMR 52, 351–364 (2012).

