## Supplemental Figures and Tables for "Multi-state structure determination and dynamics analysis reveals a new ubiquitin-recognition mechanism in yeast ubiquitin C-terminal hydrolase"

[a]Department of Chemistry, Graduate School of Science, Tokyo Metropolitan University, 1-1 Minamiosawa, Hachioji, Tokyo 192-0397, Japan; [b]International Graduate School of Arts and Sciences, Yokohama City University, 1-7-29 Suehiro-cho, Tsurumi-ku, Yokohama 230-0045; [c]RIKEN Center for Biosystems Dynamics Research, RIKEN, 1-7-22 Suehiro-cho, Tsurumi-ku, Yokohama 230-0045, Japan; [d]Molecular and Cellular Physiology Laboratory, Graduate School of Integrated Science, Yokohama City University, 1-7-29 Suehiro-cho, Tsurumi-ku, Yokohama 230-0045, Japan; [e]Graduate School of Pharmaceutical Sciences, The University of Tokyo, 7-3-1 Hongo, Bunkyo-ku, Tokyo, 113-0033, Japan; [f]Department of Medical Informatics, Research and Development Center for Medical Education, Kitasato University School of Medicine, 1-15-1 Kitasato, Minami-ku, Sagamihara, Kanagawa, 252-0374, Japan; [g]Institute of Biophysical Chemistry, Center for Biomolecular Magnetic Resonance, Goethe University Frankfurt, Max-von-Laue-Str. 9, Frankfurt am Main 60438, Germany; [h]Laboratory of Physical Chemistry, ETH Zürich, Vladimir-Prelog-Weg 2, Zürich 8093, Switzerland.

### Contents


1. **Supplementary Figure 1.** Crystal structures of YUH1 with ubiquitin aldehyde.
2. **Supplementary Figure 2.** Table summarizing the backbone and sidechain resonance assignment of YUH1.
3. **Supplementary Figure 3.** Side-chain resonance assignment of YUH1.
4. **Supplementary Figure 4.** Solution structures of YUH1 exclusively using NOEs and chemical shifts.
5. **Supplementary Figure 5.** Colormap of distance restraints from NOEs collected by the conventional structural analysis of CYANA.
6. **Supplementary Figure 6.** $T_1$, $T_2$, and $\{^1H\}$-$^{15}N$ NOE for backbone $^{15}N$ resonances of YUH1.
7. **Supplementary Figure 7.** Multiple sequence alignment of YUH1 and UCH enzymes.
8. **Supplementary Figure 8.** Cysteine mutation sites on YUH1 for the conjugation of paramagnetic tags.
9. **Supplementary Figure 9.** 2D $^1H$-$^{15}N$-HSQC spectra of cysteine mutants of YUH1.
10. **Supplementary Figure 10.** $^1H$-$^{15}N$ HSQC spectra of YUH1(N140C) tagged with DO3MA-3BrPy.
11. **Supplementary Figure 11.** Colormap of distance restraints derived from NOEs based on the 3D structural information of the multi-state ensemble conformations.
12. **Supplementary Figure 12.** SDS-PAGE analysis of the V9P mutant.
13. **Supplementary Figure 13.** 2D $^1H$-$^{15}N$ TROSY spectrum of 0.85 mM $^2H$, $^{15}N$-ubiquitin with 1.45 mM unlabeled YUH1.
14. **Table S1.** 3D NMR spectra measured for GB1 in living cells

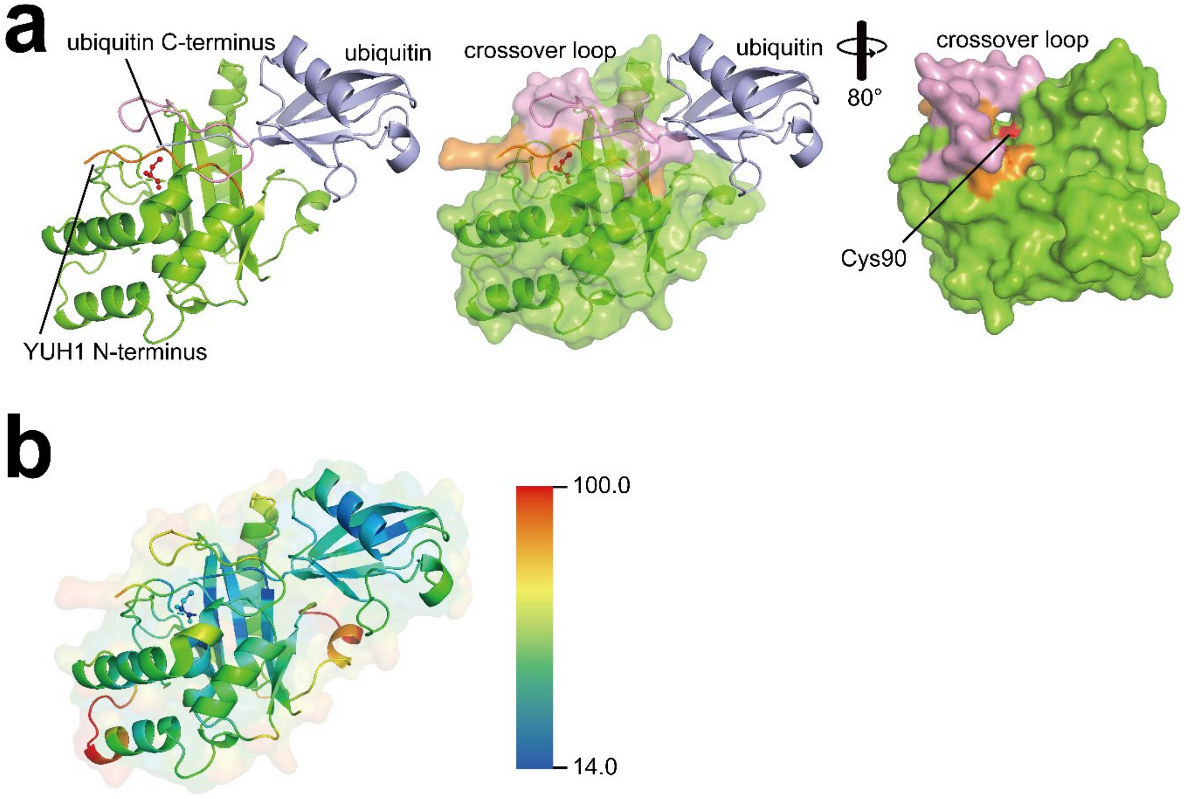


**Supplementary Figure 1. Crystal structures of YUH1 with ubiquitin aldehyde. a**, The crystal structure of the YUH1 complex with ubiquitin aldehyde (Uba1; PDBID 1CMX). It is shown as ribbon (left) and surface model (middle and right). The N-terminus, crossover loop, and Uba1 are colored in orange, pink, and light blue, respectively. Cys90 at the catalysis center is shown as ball and stick model in red. The c-terminus of Uba1 is deeply emended into the hole of the active site. **b**, Temperature factors of the crystal structure. Both ribbon and transparent surface models are rainbow colored from blue to red, progressively, from 14.0 to 100.0 of the temperature factors.

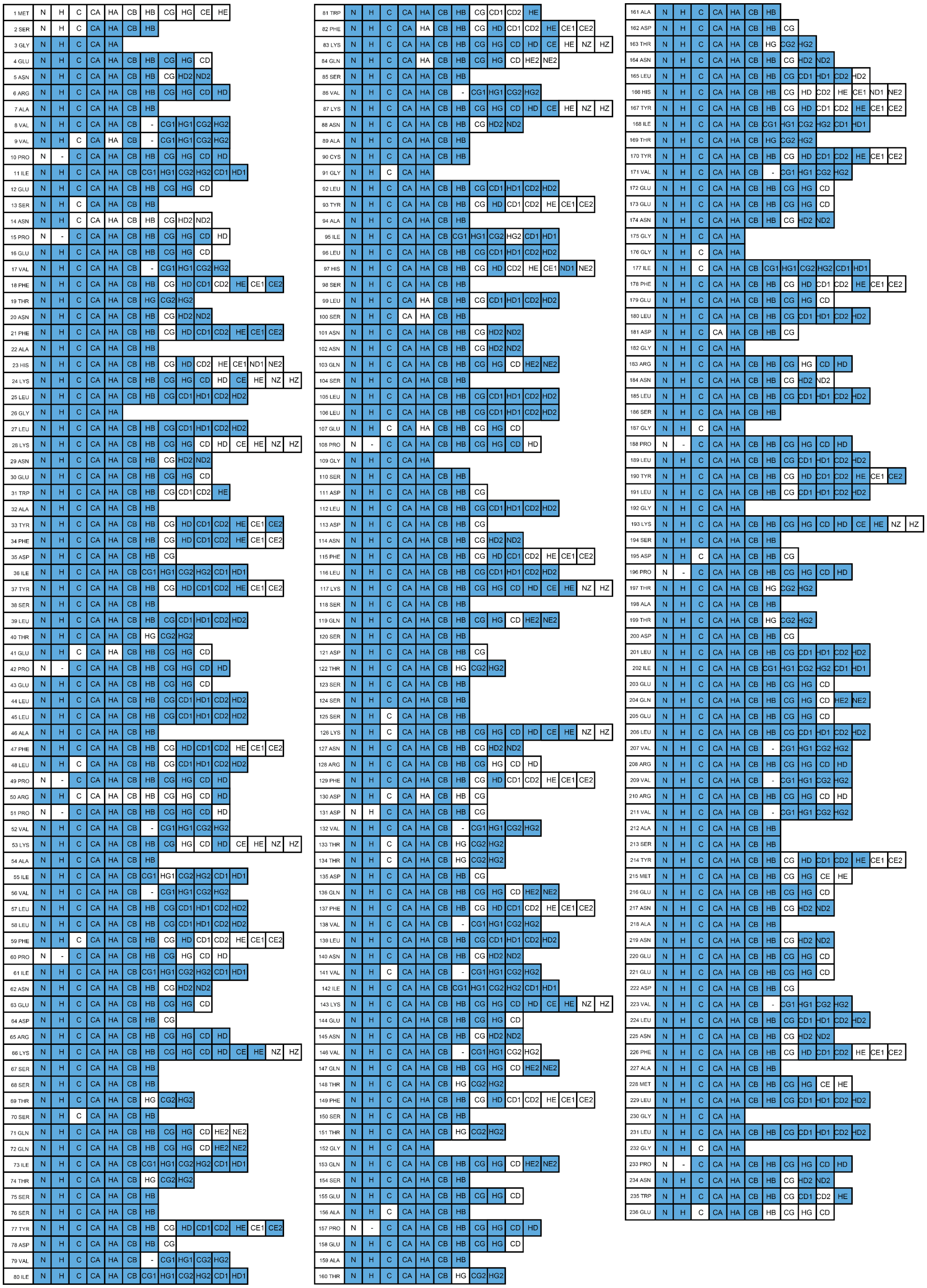

**Supplementary Figure 2. Table summarizing the backbone and sidechain resonance assignment of YUH1.** Blue and white columns indicate assigned and unassigned atoms, respectively.

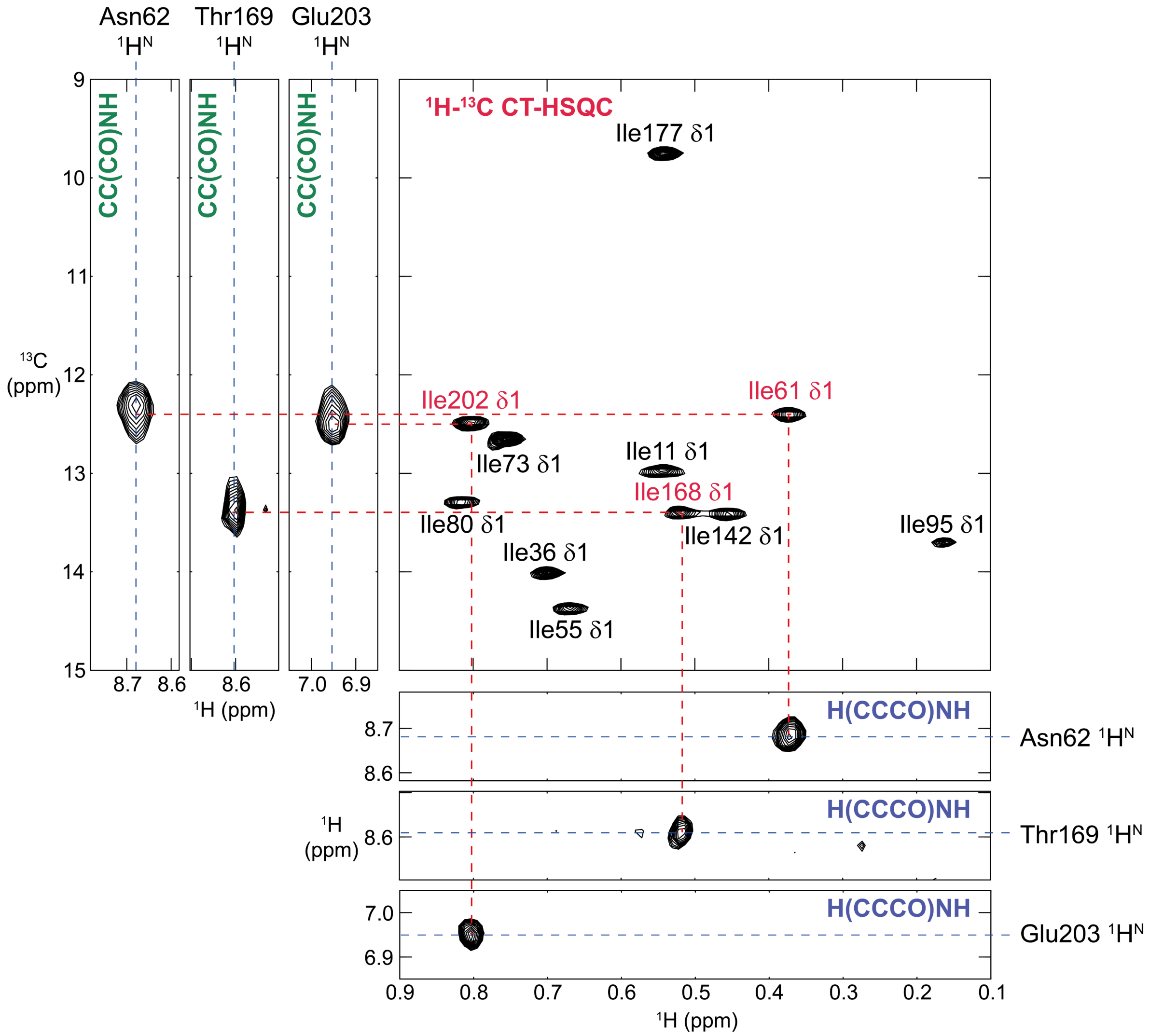


**Supplementary Figure 3. Side-chain resonance assignment of YUH1.** As examples, the manual assignment processes for the δ1 methyl $^1H/^{13}C$ resonances of Ile 61, Ile 168, and Ile 202 residues are illustrated. The $F_1(^{13}C)$-$F_3(^1H^N)$ and the $F_1(^1H)$-$F_3(^1H^N)$ slices were extracted from CC(CO)NH and H(CCCO)NH spectra, respectively, corresponding to the backbone amide $^1H^N$ and $^{15}N$ frequencies of Asn 62, Thr 169, and Glu203. From the obtained $^{13}C^{\delta1}$ and $^1H^{\delta1}$ chemical shifts, the cross peaks in 2D $^1H$-$^{13}C$ CT-HSQC were unambiguously assigned (labeled in red). The assignments for the other isoleucine δ1 resonances are shown in black.

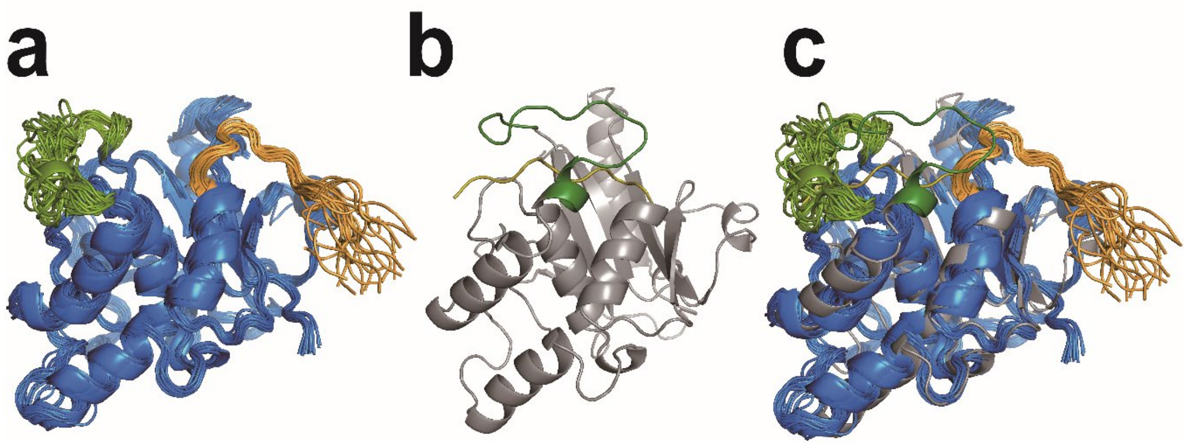


**Supplementary Figure 4. Solution structures of YUH1 exclusively using NOEs and chemical shifts. a**, Solution structures of YUH1 exclusively using NOEs and chemical shifts. **b**, the crystal structure of YUH1 from the complex (PDB ID: 1CMX). **c**, Superposition of the structures from a and b. The crossover loop and the N-terminus are colored in green and orange, respectively.

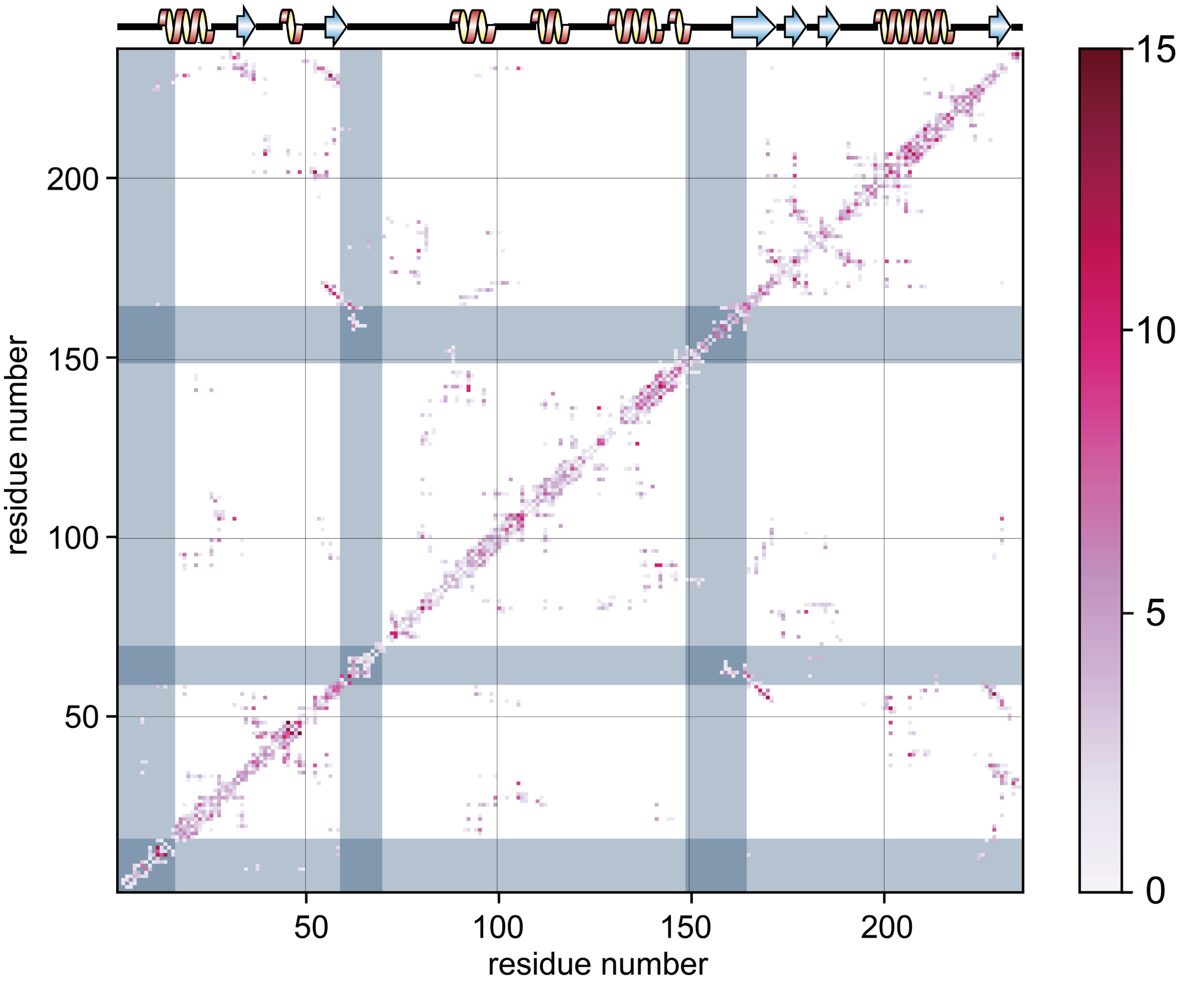


**Supplementary Figure 5. Colormap of distance restraints from NOEs collected by the conventional structural analysis of CYANA.** The grey background regions correspond to the N-terminus, and L5 and L9 loops.

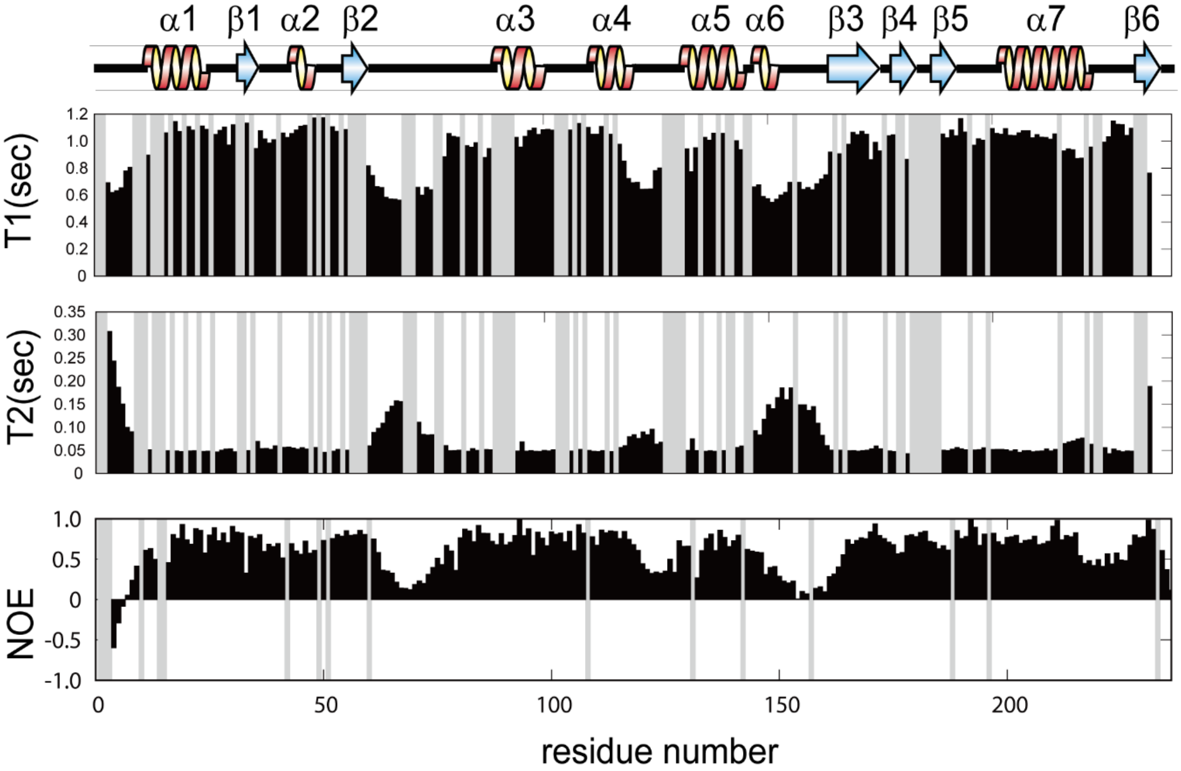


**Supplementary Figure 6.** ***T*₁, *T*₂, and {¹H}-¹⁵N NOE for backbone ¹⁵N resonances of YUH1.** Residues without data are shown in grey.

```
          N-terminal   α1        L1   β1    L2  α2   L3    β2
YUH1      MSGENRAVVPIESNPEVFTNFAHKLGLKNEWAYFDIYSLTEPELLAFLPRPVKAIVLLFP   60
UCHL1     -----MQLKPMEINPEMLNKVLSRLGVAGQWRFVDVLGL-EEESLGSVPAPACALLLLFP   54
UCHL3     --MEGQRWLPLEANPEVTNQFLKQLGLHPNWQFVDVYGM-DPELLSMVPRPVCAVLLLFP   57
UCHL3PF   -MAKNDIWTPLESNPDSLYLYSCKLGQ-SKLKFVDIYGF-NNDLLDMIPQPVQAVIFLYP   57
UCHL5     MTGNAGEWCLMESDPGVFTELIKGFGC-RGAQVEEIWSL-EPENFEK-LKPVHGLIFLFK   57

          L4     L5                 L6          α3          L7
YUH1      INEDRKSSTSQQI--------TSSYDVIWFKQSVKNACGLYAILHSLSNNQSLL--EPGS   110
UCHL1     LTAQHENFRKKQIEEL--KGQEVSPKVYFMKQTIGNSCGTIGLIHAVANNQDKLGFEDGS   112
UCHL3     ITEKYEVFRTEEEEKIKSQGQDVTSSVYFMKQTISNACGTIGLIHAIANNKDKMHFESGS   117
UCHL3PF   VNDNIVSEN---NTNDKHNLKENFDNVWFIKQYIPNSCGTIALLHLYGNLRNKFELDKDS   114
UCHL5     WQPGEEPAGSV-------VQDSRLDTIFFAKQVINNACATQAIVSVLLNCTHQ-DVHLGE   109

             α4          L8        α5       α6          L9
YUH1      DLDNFLK-SQSDTSSSKNRFDDVTTDQFVLNVIKENVQTFSTGQSE---APEATADTNLHY   167
UCHL1     VLKQFLSETEKM-SPED-RAKCFEKNEAIQA---AHDAVAQEGQCR---V---DDKVNFHF   162
UCHL3     TLKKFLEESVSM-SPEE-RARYLENYDAIRV---THETSAHEGQTE---APSIDEKVDLHF   170
UCHL3PF   VLDDFFNKVNEM-SAEK-RGQELKNNKSIEN---LHHEFC--GQVE---NRDDILDVDTHF   165
UCHL5     TLSEFKEFSQSF-DAAM-KGLALSNSDVIRQ---VHNSFARQQMFEFDTKTSAKEEDAFHF   165

          β3 L10 β4    L11    β5      L12             α7         L13
YUH1      ITYVEENGGIFELDGRNLSGPLYLGKSDPTATDLIEQELVRVRVASYMENANEEDVLNFA   227
UCHL1     ILFNNVDGHLYELDGRMPFPVNH----GASSEDTLLKDAAK-VCREFTERE--QGEVRFS   215
UCHL3     IALVHVDGHLYELDGRKPFPINH----GETSDETLLEDAIE-VCKKFMERD--PDELRFN   223
UCHL3PF   IVFVQIEGKIIELDGRKDHPTVH----CFTNGDNFLYDTGKIIQDKFIEKC--KDDLRFS   219
UCHL5     VSYVPVNGRLYELDGLREGPIDL----GACNQDDWISAVRPVIEKRIQKYS--EGEIRFN   219

          β6  C-terminal
YUH1      MLGLGPNWE---------------------------------------------------   236
UCHL1     AVALCKAA----------------------------------------------------   223
UCHL3     AIALSAA-----------------------------------------------------   230
UCHL3PF   ALAVIPNDNFDII-----------------------------------------------   232
UCHL5     LMAIVSDRKMIYEQKIAELQRQLAEEEPMDTDQGNSMLSAIQSEVAKNQMLIEEEVQKLK   279

YUH1      --------------------------------------------------   236
UCHL1     --------------------------------------------------   223
UCHL3     --------------------------------------------------   230
UCHL3PF   --------------------------------------------------   232
UCHL5     RYKIENIRRKHNYLPFIMELLKTLAEHQQLIPLVEKAKEKQNAKKAQETK   329
```

**Supplementary Figure 7. Multiple sequence alignment of YUH1 and UCH enzymes.**
The green boxes show identical residues with YUH1.

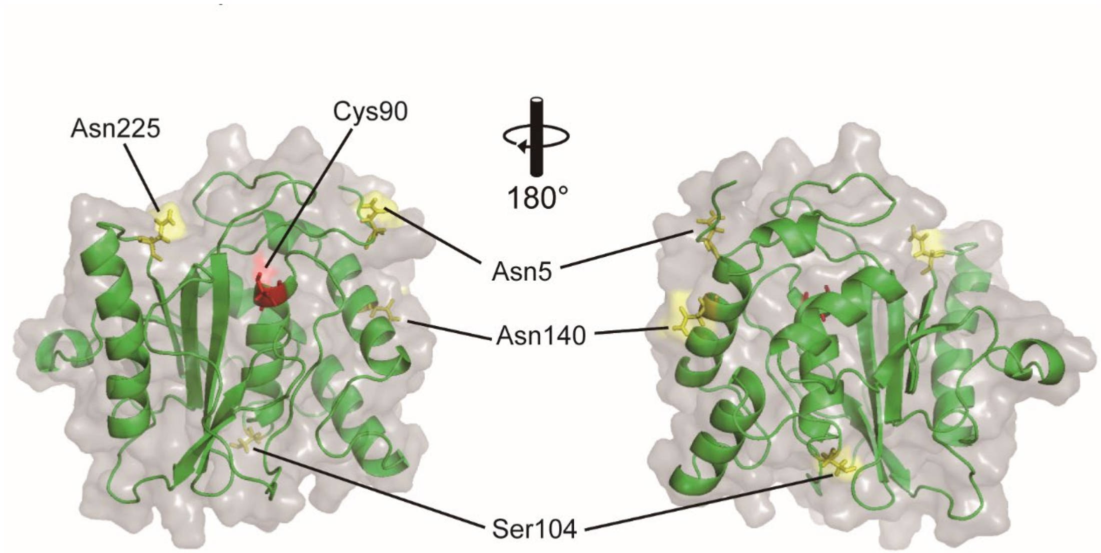


**Supplementary Figure 8. Cysteine mutation sites on YUH1 for the conjugation of paramagnetic tags.** The residues replaced by cysteines and the catalytic site (Cys90) are color-coded in yellow and red, respectively.

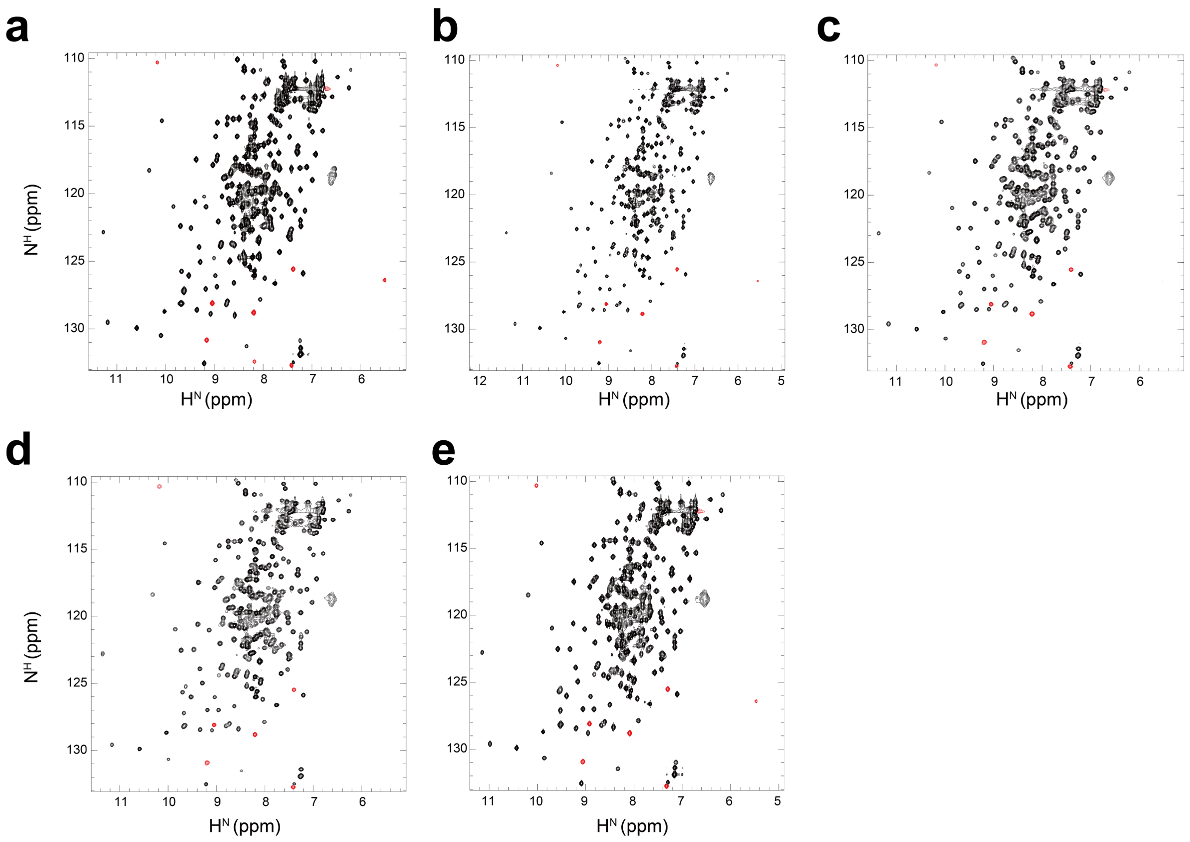


**Supplementary Figure 9. 2D $^{1}$H-$^{15}$N-HSQC spectra of cysteine mutants of YUH1. a,** wild type **b,** C90S/N5C mutant **c**, C90S/S104C mutant **d**, C90S/N140C mutant **e**, C90S/N225C mutant. In all NMR spectra, positive and negative (aliased on the $^{15}$N axis) signals are color-coded in black and red, respectively.

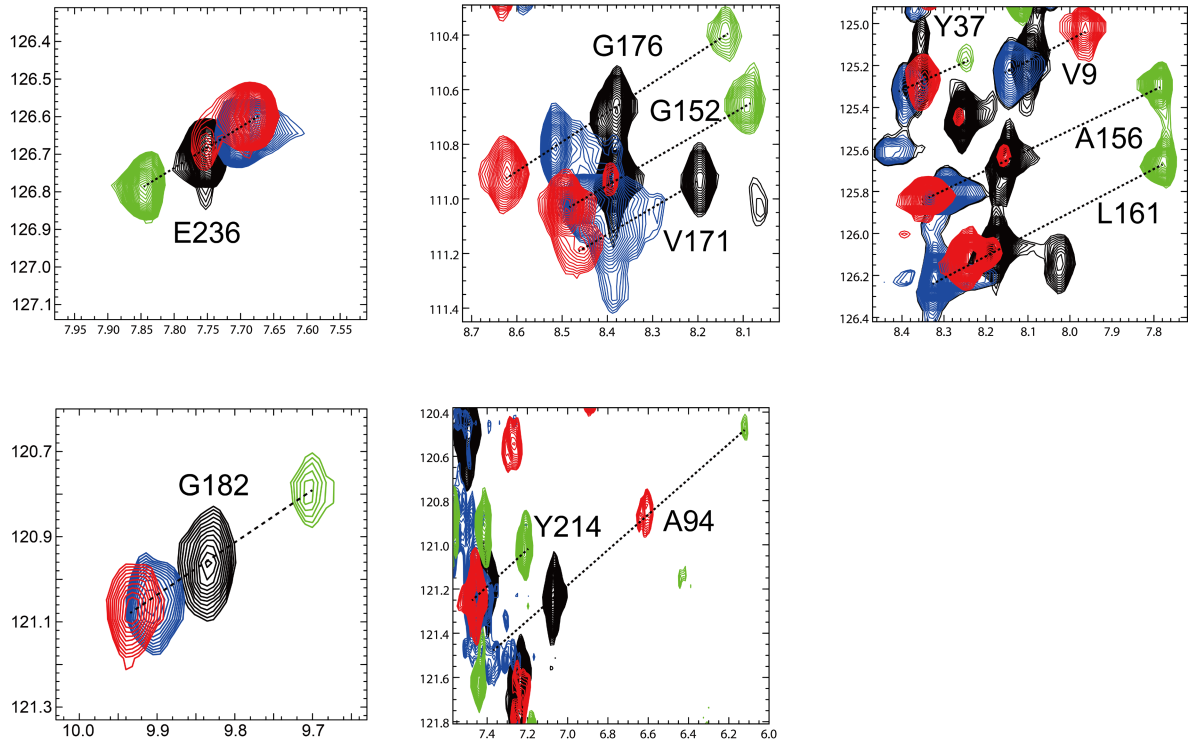


**Supplementary Figure 10. $^1H$-$^{15}N$ HSQC spectra of YUH1(N140C) tagged with DO3MA-3BrPy.** Several representative cross peaks from Figure 2c are shown separately. Diamagnetic $Lu^{3+}$ and paramagnetic $Dy^{3+}$, $Tb^{3+}$, and $Tm^{3+}$ are colored in black, red, blue, and green, respectively.

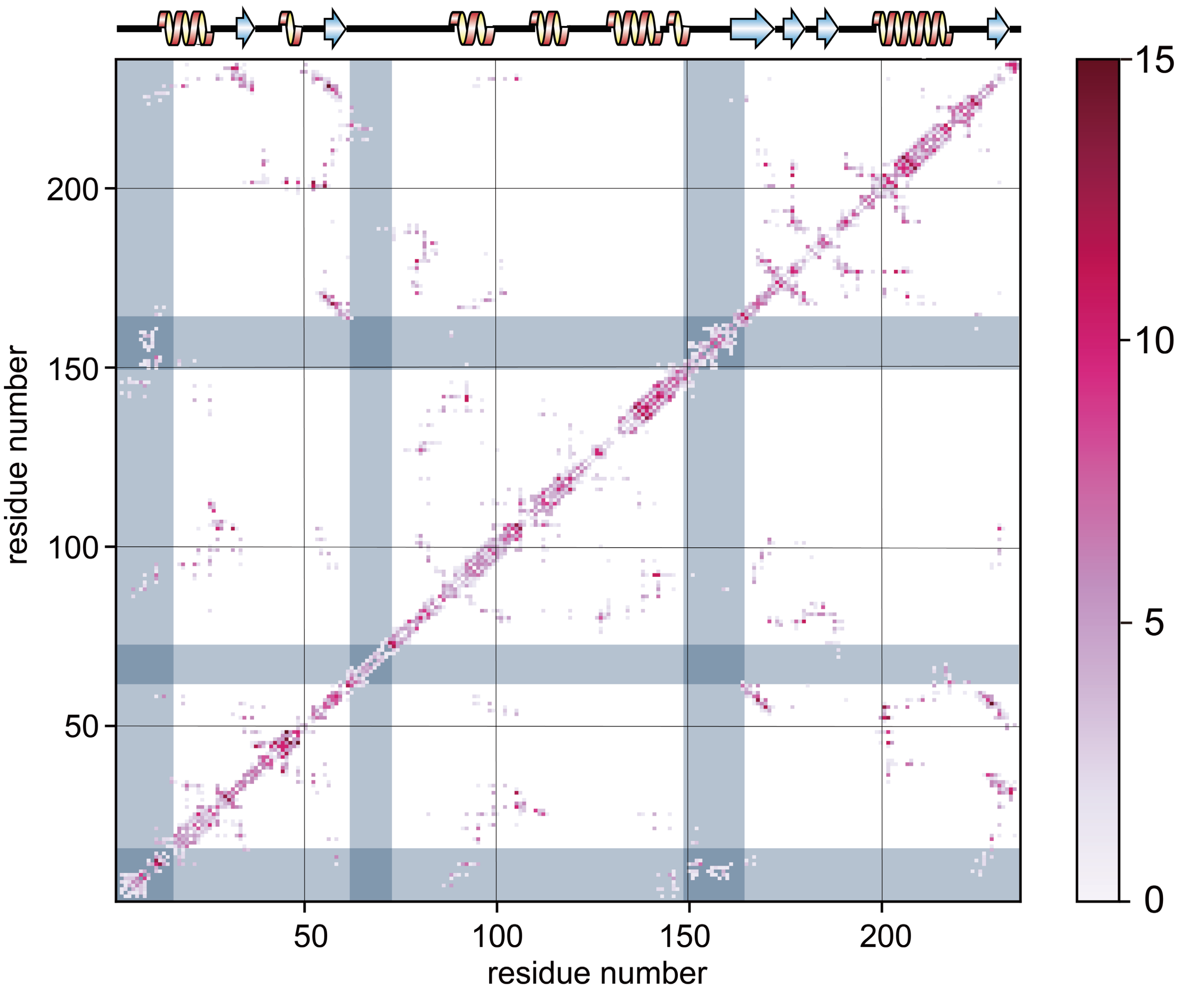


**Supplementary Figure 11. Colormap of distance restraints derived from NOEs based on the 3D structural information of the multi-state ensemble conformations.** The grey background regions correspond to the N-terminus, and L5 and L9 loops.

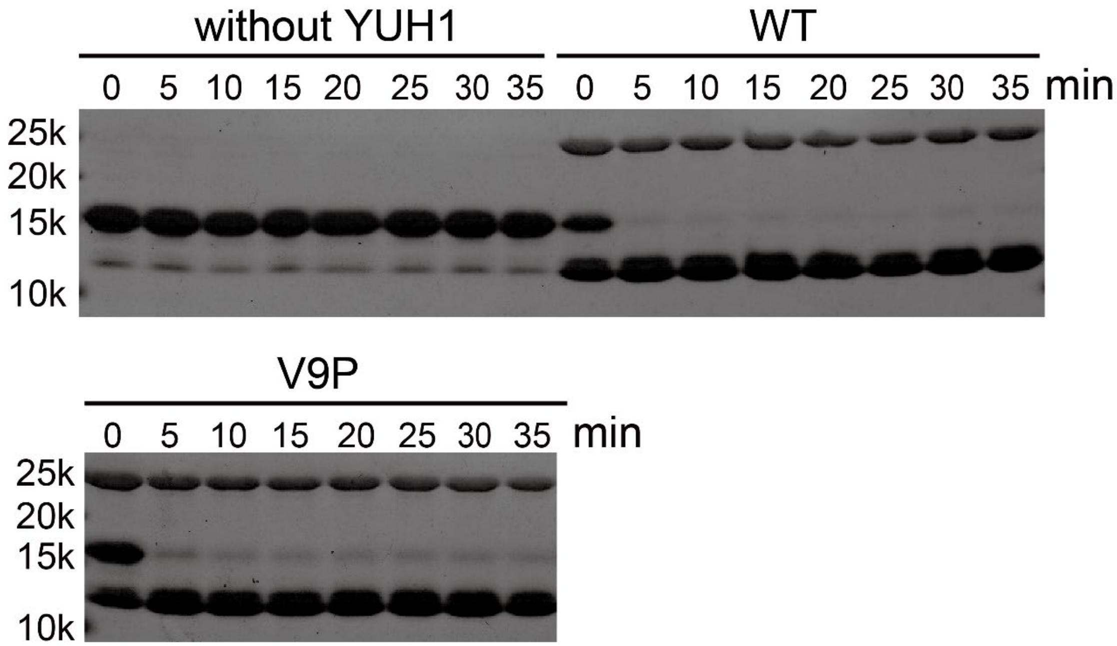


**Supplementary Figure 12. SDS-PAGE analysis of the V9P mutant.** YUH1 hydrolysis activity of Ub-Rec33 fusion protein. Reaction samples were collected at the given time points and subjected to SDS-PAGE electrophoresis without (top right), with (top left) wild type YUH1, and with the V9P mutant (bottom). The reactions shown here were performed at 25 °C. YUH1, Ub-Rec33 fusion protein, and cleaved Ub (in addition to a $His_6$ tag at the N-terminus) appear as a band at about 26, 15, and 11 kDa, respectively. Cleaved Rec33 is invisible on these electrophoresis gels because of its small size of 4 kDa.

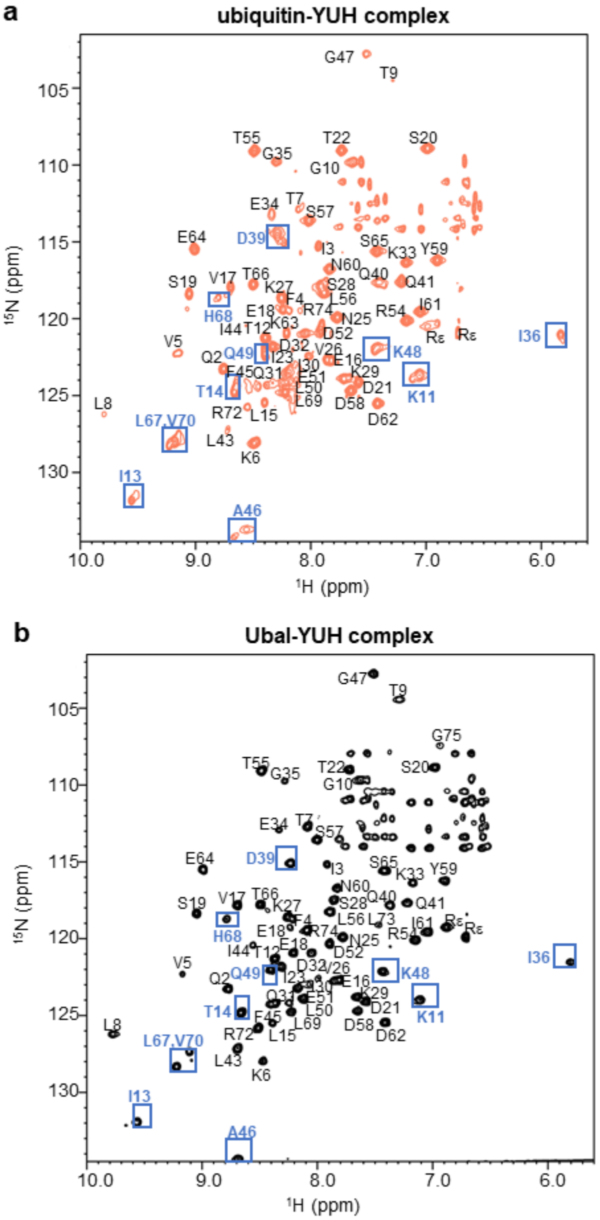


**Supplementary Figure 13. 2D $^{1}$H-$^{15}$N TROSY spectrum of 0.85 mM $^{2}$H, $^{15}$N-ubiquitin with 1.45 mM unlabeled YUH1 (a) and 0.5 mM [$^{2}$H, $^{15}$N]Ubal-unlabeled YUH1 complex (b).** Assignments of each signal are indicated. Resonances from residues that exhibited two signals in (a) are enclosed in boxes.

**Supplementary Table 1. YUH1 structure statistics**

| Quantity[a] | conventional method | multi-state calculation |
|---|---|---|
| Assigned $^1H/^{13}C/^{15}N$ chemical shifts | 2374/1176/304 | 2374/1176/304 |
| NOE distance restraints[b] | 1164/979/1462 | 1541/1340/1917 |
| Dihedral angle restraints ($\phi/\psi$) | 288 | 288 |
| Max. distance restraint violation (Å) | 0.15 ± 0.02 | 0.09 ± 0.00 |
| Max. dihedral angle restraint violation (º) | 4.89 ± 0.80 | 5.56 ± 4.07 |
| Deviations from idealized geometry: | | |
| Bond lengths (Å) | 0.0142±0.0001 | 0.0147±0.001 |
| Bond angles (º) | 2.08 ± 0.03 | 1.14 ± 0.01 |
| AMBER energy (kcal/mol) | – 8,818 ± 133 | – 9,174 ± 171 |
| AMBER vdW energy (kcal/mol) | – 627 ± 21 | – 819 ± 24 |
| Ramachandran plot statistics[c] (%) | 78/18/3/0 | 84/14/2/0 |
| Backbone RMSD (Å)[d] | 0.59 ± 0.08 | 0.77 ± 0.06 |
| All heavy atom RMSD (Å)[d] | 1.05 ± 0.07 | 1.45 ± 0.07 |
| Backbone RMSD to the reference (Å)[e] | 6.16 | 1.31 |
| All heavy atom RMSD to the reference (Å)[e] | 6.62 | 2.03 |

[a]Where applicable, the average value and the standard deviation over the 20 energy-refined conformers. For the conventional method, 20 conformers with the lowest target function were used for the statistics. For the multi-state calculation, four conformers were randomly selected from each of the five clusters (total 20 conformers).

[b]Short/medium/long-range distance restraints derived from NOESY spectra. Percentage of residues in the most favored/additionally allowed/generously allowed/disallowed regions of the Ramachandran plot according the program PROCHECK.

[d]RMSD to the mean structure for residues 15–62, 78–148, and 165–236

[e]RMSD between the closest structure to the reference among the ensemble conformations and the reference structure (the crystal structure of the YUH1-Ubal complex).
